# DRAG *in situ* barcoding reveals an increased number of HSPCs contributing to myelopoiesis with age

**DOI:** 10.1101/2022.12.06.519273

**Authors:** Jos Urbanus, Jason Cosgrove, Joost Beltman, Yuval Elhanati, Rafael de Andrade Moral, Cecile Conrad, Jeroen W van Heijst, Emilie Tubeuf, Arno Velds, Lianne Kok, Candice Merle, Jens P Magnusson, Jonas Frisén, Silvia Fre, Aleksandra M Walczak, Thierry Mora, Heinz Jacobs, Ton N. Schumacher, Leïla Perié

## Abstract

Ageing is associated with changes in the cellular composition of the immune system. During ageing, hematopoietic stem and progenitor cells (HSPCs) that produce immune cells are thought to decline in their regenerative capacity. However, HSPC function has been mostly assessed using transplantation assays, and it remains unclear how HSPCs age in the native bone marrow niche. To address this issue, we developed a novel in situ single cell lineage tracing technology to quantify the clonal composition and cell production of single cells in their native niche. Our results demonstrate that a pool of HSPCs with unequal output maintains myelopoiesis through overlapping waves of cell production throughout adult life. During ageing, the increased frequency of myeloid cells is explained by greater numbers of HSPCs contributing to myelopoiesis, rather than increased myeloid output of individual HSPCs. Strikingly, the myeloid output of HSPCs remained constant over time despite accumulating significant transcriptomic changes throughout adulthood. Together, these results show that, unlike emergency myelopoiesis post-transplantation, aged HSPCs in their native microenvironment do not functionally decline in their regenerative capacity.

## Introduction

Immune cells are constantly replenished throughout an organism’s lifetime by hematopoietic stem cells (HSCs). While this replenishment is particularly important for short lived immune cells such as granulocytes and monocytes, longer lived immune cells such as lymphocytes also require input from hematopoiesis in addition to homeostatic proliferation. It has previously been demonstrated that during ageing blood cell production shifts toward myeloid cells at the expense of lymphoid cells ^1^, a change that correlates with a higher risk of several myeloid-associated pathologies, including myelodysplastic syndromes and leukemia.

Within the murine bone marrow, age-related changes in myeloid cell numbers are accompanied by an increase in immunophenotypic stem cell (HSC) frequency or numbers ^1–5^. Downstream of HSCs, lymphoid biased MPPs descrease in frequency ^6^, followed by a decreased frequency in downstream common lymphoid progenitors ^4^ and an increased frequency of granulo-monocyte progenitors ^4^. At the cellular level, the increase in myeloid production may happen through two non-exclusive mechanisms: an increase in the number of myeloid-biased hematopoietic stem and progenitor cells (HSPCs), or an increase in the number of myeloid cells that are produced per individual HSPC. The first mechanism has been well documented post-transplantation ^6–10^ whereas the second mechanism remains controversial ^5, 7, 9, 11^. At the molecular level, aged HSPCs upregulate stress response and inflammation related gene signatures, as well as genes involved in myeloid differentiation ^4, 5, 12, 13^. Aged HSCs show increased expression of a self-renewal related gene expression program ^12, 14^, while aged HSPCs increase expression of differentiation-related programs ^12, 14^ relative to young HSPCs. These molecular changes, together with the decreased rate of cell production of old HSCs in competitive transplantation with young HSCs^1, 15^, the increased number of myeloid-biased HSCs^3, 6, 7, 15, 16^ and the decreased self-renewal of HSC after secondary transplantation^3, 8, 15^ has led to a model in which aged HSPC exhaustion is a hallmark of an ageing immune system^17^.

Importantly, consolidating data from native and post-transplantation hematopoiesis is non-trivial, with recent reports highlighting important differences between them^18^. Using *in situ* barcoding, Camargo and colleagues demonstrated that in young adult mice a larger number of HSCs contributes to steady state hematopoiesis as compared to post-transplantation hematopoiesis ^19^. In addition, fate mapping studies have shown that the dynamics of HSC activation in native hematopoiesis ^20^ differ from those observed after transplantation ^21^. Given that most functional measurements of aged HSPCs come from transplantation assays, further work on the functional characterization of HSPCs within the native bone marrow microenvironment is required. One study using confetti mice showed a decrease in HSC clonal diversity with age ^22^ but the modest diversity of this system is insufficient to uniquely label each cell within the HSC pool, estimated to comprise 17,000 cells in a single mouse ^23^. Furthermore, HSPCs have been shown to display functional heterogeneity with respect to self-renewal, differentiation and proliferation capacity ^24–26^ and the definition of HSPCs has evolved recently to include several new, functionally distinct, murine MPP subsets ^27–29^. In summary, existing models of HSPC ageing are based on transplantation studies or population-level assays. However, recent studies have identified important differences between native and transplantation hematopoiesis, highlighting the need for single-cell resolution assays to resolve HSPC heterogeneity. *In situ* barcoding approaches can address these two key limitations, and may thereby improve our understanding of the cellular dynamics that drive ageing of the immune system.

To elucidate the effect of ageing on HSPCs in their natural niche at single-cell resolution, we developed the DRAG mouse, a novel *in situ* single cell lineage tracing technology that exploits the process of VDJ recombination. DRAG barcoding revealed that steady state adult myelopoiesis is sustained by overlapping waves of cell production throughout adulthood. Furthermore, the increased rate of myeloid cell production during ageing is explained through an increase in the number of myeloid producing HSPCs, rather than an increase in the number of myeloid cells produced per individual HSPC. Single-cell RNA sequencing analysis of HSPCs across adulthood is consistent with this model, suggesting a reduced frequency of quiescent stem cells, and the emergence of age-associated active progenitor subsets. Collectively, our data reveal that, in the native bone marrow microenvironment, individual aged HSPCs produce myeloid cells at the same rate as young HSPCs, despite the accumulation of transcriptomic changes associated with stress and inflammation. These data provide evidence that HSPC in their native niche are not exhausted in their capacity to produce myeloid cells.

## Results

### A novel quantitative in situ barcoding system

Taking advantage of the capacity of the VDJ recombination system to produce a high degree of genetic diversity in the lymphoid lineages, we designed a DNA cassette, termed DRAG (Diversity through RAG), with the aim to allow endogenous barcoding in an organism in a temporally controlled manner (Fig. 1A). The DRAG system has been designed such that upon CRE induction, a segment between two loxP sites is inverted, leading to the expression of both the RAG1 and 2 enzymes and Terminal deoxynucleotidyl transferase (TdT). Such expression then leads to the semi-random RAG-mediated recombination of synthetic V-, D- and J-segments, with additional diversity being generated both by nucleotide deletion and TdT-mediated N-addition at the junction sites. Notably, as the RAG/TdT cassette and recombination signal sequences (RSSs) are spliced out during this recombination step, further recombination of the DRAG locus is prevented, and any generated VDJ sequence is thus stable over time. Finally, recombination of the DRAG locus results in the removal of a BGH polyA site that precludes GFP expression in the DRAG configuration before recombination, allowing one to identify barcode^+^ cells by flow cytometry or imaging (Fig. 1A).

**Figure 1:**
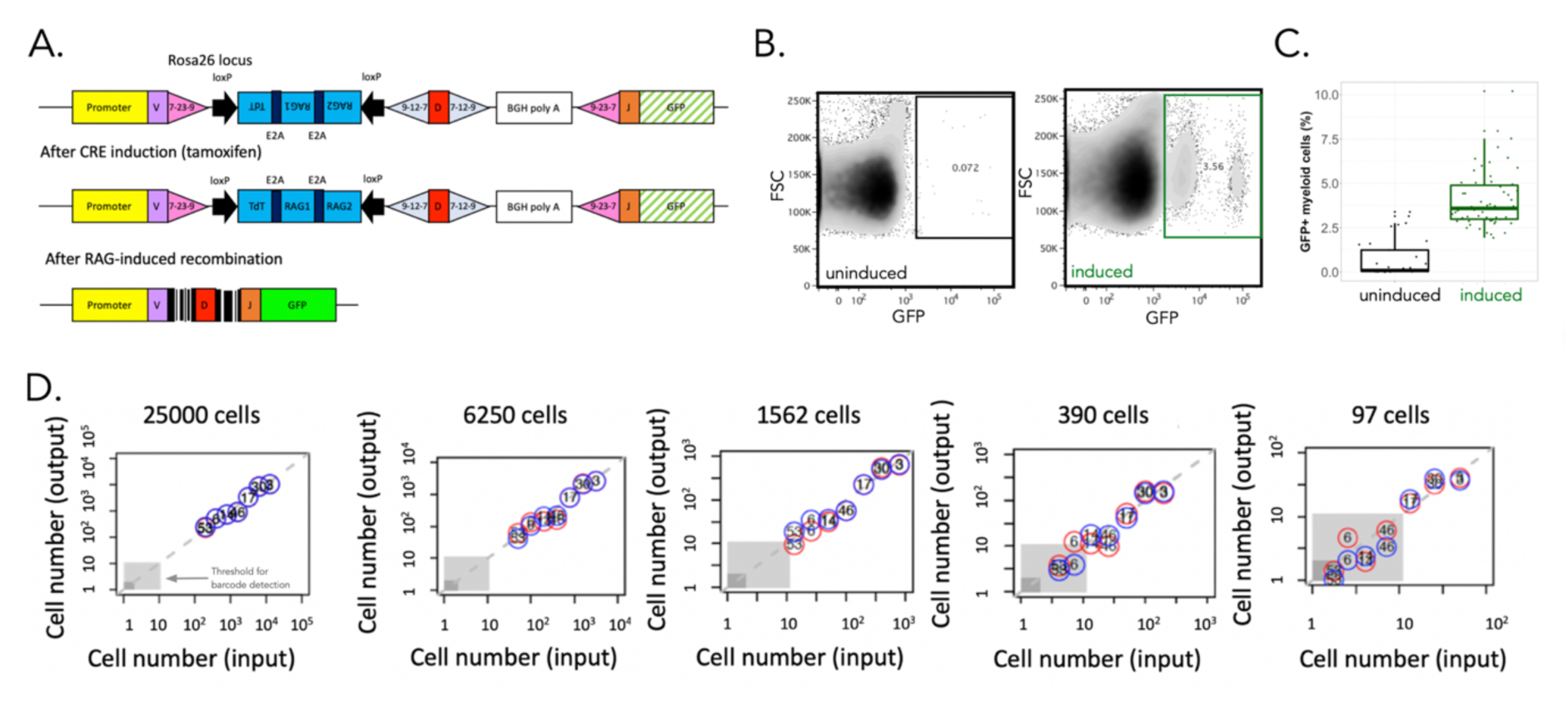
A quantitative DRAG in situ barcoding system. A. Description of the DRAG cassette, as inserted into the Rosa 26 locus before and after induction. DRAG recombination is induced by Cre activity and resulting barcode sequences are used for lineage tracing. B. Example of GFP expression in myeloid cells (CD11b^+^ CD19^−^ CD3^−^ CD11c^−^) in blood 6.5 months after tamoxifen (induced) or vehicle (control) administration. Within the GFP positive gate, a GFP^mid^ and GFP^high^ population is observed in myeloid cells. Both populations contain successfully recombined barcodes, and heterogeneity in GFP marker expression is likely due to the labelling of heterogeneous cell types with the pan myeloid marker cd11b (Figure S5A-B). In line with this, such heterogeneous GFP expression was not observed in non-myeloid cells. C. Percentage of GFP^+^ myeloid cells of total myeloid cells in tamoxifen-induced (green) and control (black) (n=5 and n=3 mice respectively, all sampled over 13 months). Median and interquartile range with whiskers extending to the minimum and maximum values. D. False positive rate and sensitivity of barcode detection. 7 MEF clones with known DRAG barcodes, were mixed in different numbers, and the input cell numbers of all MEF clones were compared to experimentally determined numbers upon PCR, sequencing and analysis. Number in circles correspond to MEF clone numbers, Red and blue circles indicate technical replicates. Grey area indicates lower thresholds for barcode detection as used during data processing.

To quantify the sensitivity, specificity and fidelity of the DRAG barcoding system, we benchmarked the DRAG barcoding system *in vitro* and *in vivo*. First, to understand DRAG recombination patterns *in vitro*, we isolated embryonic fibroblasts (MEF cells) from CAGCre-ER^+/−^ DRAG^+/−^ mice, induced DRAG recombination with tamoxifen, and derived MEF clones (n=24) that carried a single DRAG barcode by limited dilution. Recombined DRAG loci were characterized by insertions and deletions between the VDJ segments (Table S1), and only 2 of the 24 barcodes were shared between MEF clones. Longitudinal analysis of barcode sequences by Sanger and deep sequencing demonstrated that recombined sequences were stable over time in all clones tested (Table S2). To allow robust detection, identification and quantification of DRAG barcodes, we developed a processing platform that incorporates unique molecular identifiers (UMI) during PCR amplification (Fig. S1B Sup Method) and a strict filtering pipeline of sequencing data. Application of this strategy to defined mixtures of 7 MEF clones that each contain a different barcode demonstrated that this platform allows the robust detection and quantification of barcodes (Fig. 1D). With regards to barcode specificity, in these mixtures, the number of false positive DRAG barcode events was 0 over 14, except for one mixture in which we had 1 false positive over 14. With regards to sensitivity of detection, clones equal or greater than 10 cells were efficiently identified in samples of as few as 97 cells (Fig. 1D); and clones equal or greater than 100 cells were identified in samples of 25,000 cells (Fig. 1D), corresponding to as little as 0.4% of the total cell population. Furthermore, across the entire detection range, experimentally observed frequencies were highly correlated to input frequencies (Fig. 1D), demonstrating that DRAG permits quantitative analyses of clonal output.

To characterize the DRAG barcoding system *in vivo,* we induced barcode recombination by tamoxifen administration in CAGCre-ER^+/−^ DRAG^+/−^ mice and analysed hematopoietic cells from blood samples taken 6 months later. Induction of Cre resulted in a 5- to 6-fold increase in barcode-labeled (GFP^+^) myeloid cells (Fig. 1B-C), relative to mock-induced DRAG mice. In addition, no variation was observed in the percentage of myeloid and lymphoid cells produced from recombined (GFP^+^) or unrecombined (GFP^−^) cells (Fig. S1A), indicating that DRAG induction appears neutral with respect to hematopoietic cell differentiation. Deep sequencing analysis of barcode sequences processed 15 months post Cre-induction confirmed the high diversity of the DRAG barcoding system *in vivo*. Specifically, barcodes were characterized by up to 15 nucleotide insertions and 32 nucleotide deletions (Fig. S2A-B), and D-segment inversion was observed in 12% of cases. Lastly, to assess the likelihood that the same DRAG recombination pattern occurs independently in 2 or more cells *in vivo* we applied a mathematical model^30^ to infer the probability that a given barcode will be produced in the DRAG system (P_gen_) using experimental data as input (Suppl. methods and Table S3–8). This experimentally validated approach (Fig. S2, Suppl. Methods) enabled us to identify sequences with a high generation probability (Fig. S2C), such that they can be filtered from the data prior to downstream analyses. On the basis of these results, we selected a P_gen_ estimate of <10^−4^ for further data analysis (Fig. S2D-E). At this probability cut-off, 61% of barcode sequences were retained, and 92% of these retained barcodes were unique to an individual mouse, yielding hundreds of barcodes that could be used for analysis.

Together, these data show that the DRAG mouse model allows the generation for high barcode diversity *in vivo* without a requirement for cell transplantation, and that the prevalence of these barcodes in downstream progeny can be detected in a quantitative manner.

### DRAG labeling of hematopoietic stem and progenitor cells

Having established the feasibility of in vivo barcoding in DRAG mice, we applied the system to study native hematopoiesis. Given the ubiquitous expression of the CAGCre-ER^TM^ driver, tamoxifen-based induction will result in the labeling of both HSPCs, but also committed progenitors and differentiated cells. Importantly, at late time points (month) after induction, turnover of short-lived committed progenitors and differentiated cells, and replacement by progeny of long-lived cells, has occurred (Fig. 2A). Thus, the barcodes observed several months after induction will be inherited from ancestor cells that qualify as long-term repopulating cells *in vivo* ^31^. Importantly, this unbiased functional definition of long-term output towards a short-lived downstream cell population is independent of surface markers or HSC-selective gene promoters to drive Cre expression. To directly test whether the DRAG system homogeneously labels the HSPC compartment, we performed 10X single cell RNA-sequencing on both GFP^+^ and GFP^−^ bone marrow HSPCs. After data QC and processing (Suppl. methods), unsupervised Louvain clustering resulted in 8 clusters (Fig. S3A-C) that were annotated by mapping to the previously described gene expression signatures of long-term HSCs and multipotent progenitors^28, 32^ (Fig. 2B and C). GFP^+^ and GFP^−^ HSPC cells were distributed equally among the clusters (Fig. 2D and E), with the exception of MPP3 that was enriched in GFP^+^ cells (Fishers exact test p < 0.001), but this effect was not statistically significant when the frequency of GFP labelling was assessed by flow cytometry (Fig S3D). Importantly, GFP^+^, and hence barcode-labeled, cells were also observed in the long-term HSC associated clusters, characterized by high expression of the long-term HSC gene signature ^32^ (Fig. 2C), high expression of *Ly6a,* and low expression of *Cd48* (Fig. S3E). Thus, the DRAG system efficiently labels the HSPC compartment, including long-term hematopoietic stem cells.

**Figure 2:**
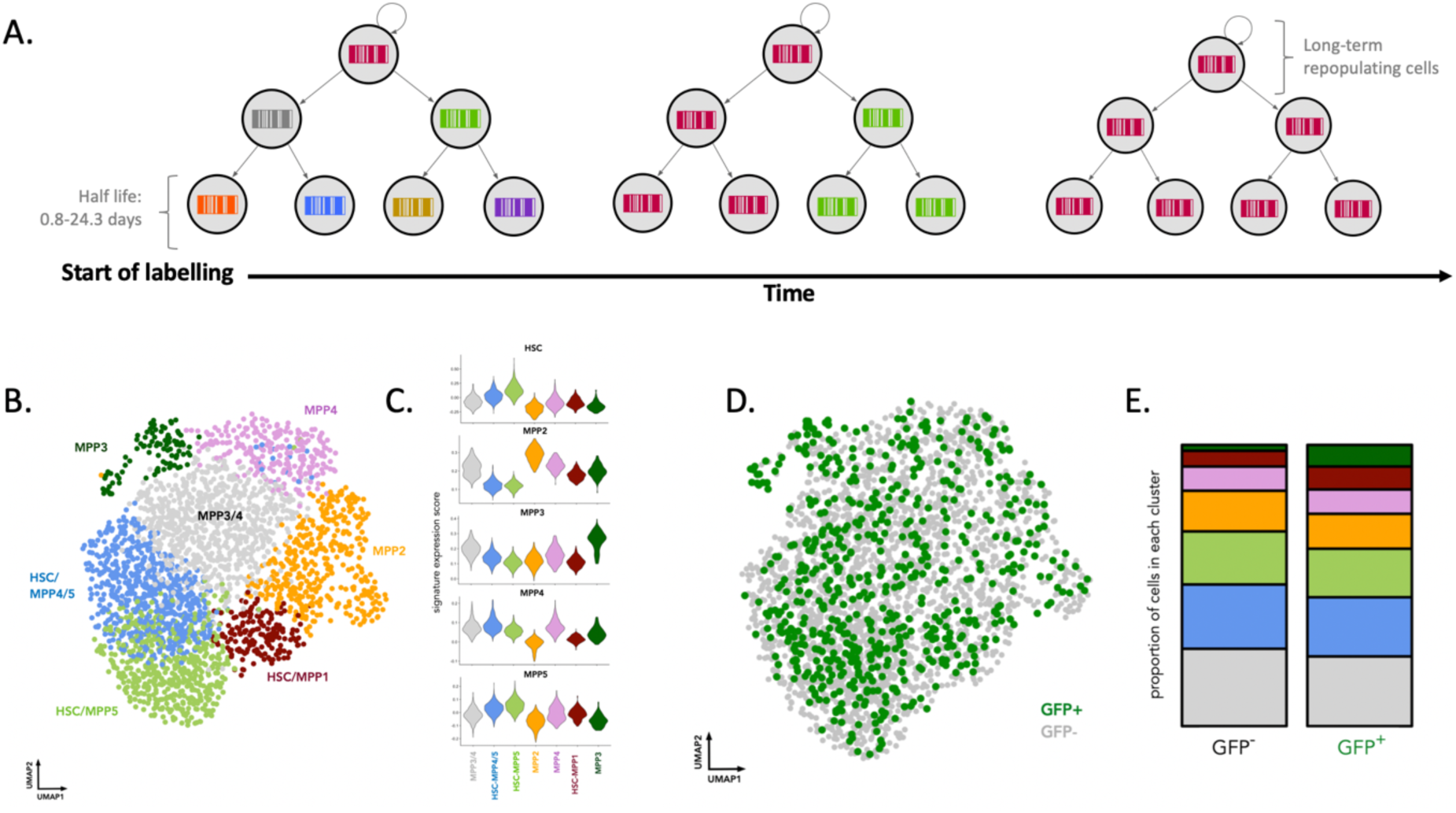
Identity of DRAG barcode labelled HSPC cells. A. At the start of DRAG labeling, tamoxifen-induced Cre-ER^TM^ activity will yield DRAG barcodes in stem cells, progenitor cells, and downstream differentiated cells. At later time points (month) after induction, turnover of short-lived committed progenitors and differentiated cells, and replacement by progeny of long-lived cells has occurred, and DRAG barcodes observed in short-lived differentiated cell pools are derived from long-term repopulating cells. B. Six months post tamoxifen induction, HSPC cells (LSK: sca1^+^ckit^+^ cells) GFP^+^ and GFP^−^ were scRNA sequenced using the 10X 3’end protocol (data from 2 mice induced at 20 weeks). UMAP representation of the data, with key subpopulations obtained by Louvain clustering highlighted. C. Published gene expression signatures ^27^ were used to annotate and quantify the clusters in B. D. Distribution of GFP^+^ cells throughout the UMAP embedding of the data E. The proportion of cells in each cluster from B within either GFP^+^ or GFP^−^ cells.

### Steady state myelopoiesis is maintained by overlapping waves of cell production

To study the clonal dynamics of myelopoiesis at steady state, we analysed the distribution of barcodes across different developmental compartments of the bone marrow. Specifically, bone marrow HSPCs (LSK: sca1^+^ckit^+^), myeloid progenitors (MP: myeloid progenitors, sca1^−^ckit^+^) and myeloid cells (CD11b^+^) were isolated at 15-month post-induction in a cohort of DRAG mice that received tamoxifen induction between 6-14 weeks (Fig. 3A and S4A-B). Following initial filtering, quality checks and removal of frequently occurring barcodes, correlations between barcode abundance in duplicate samples (LSK: 0.5 +/−0.1, MP: 0.7 +/− 0.1, M: 0.6 +/− 0.2) were calculated to assess the consistency between technical replicates. Following this quality control steps, barcode sharing analysis revealed a number of different fates with only 13.7% (+/−5% SD between mice) of barcodes shared across HSPCs, myeloid progenitors and myeloid cells. These multi-outcome clones were among the most prolific, producing 41.4% (+/− 12.2% SD between mice) of myeloid cells, and representing 69.9% (+/−23.7% SD between mice) of HSPCs (Fig. 3C and D). Interestingly, a number of barcodes detected in HSPCs were not detected in downstream developmental compartments, suggesting that their contribution to myelopoiesis at this timepoint was limited. These HSPC-restricted barcodes produced 24.6% (+/− 19% SD between mice) of the total HSPCs (Fig. 3C and D). In addition, we detected barcodes that were abundant in myeloid progenitors and myeloid cells but below the threshold of detection in HSPCs. These MP-M restricted barcodes were producing 48% (+/−23% SD between mice) of myeloid progenitors and 43.2% (+/−8% SD between mice) of myeloid cells (Fig. 3C and D). Notably, while MP-M and HSPC-MP restricted barcodes were both observed, no barcodes were detected in both HSPC and myeloid cells without being detected in myeloid progenitors (HSPC-M class, Fig. 3C), arguing against stochastic detection of DRAG barcodes as a major confounder. Importantly, barcode outcomes were independent of barcode generation probability, suggesting that the barcode patterns we observed across developmental compartments were not due to limitations in detection sensitivity (Fig. S4F).

**Figure 3:**
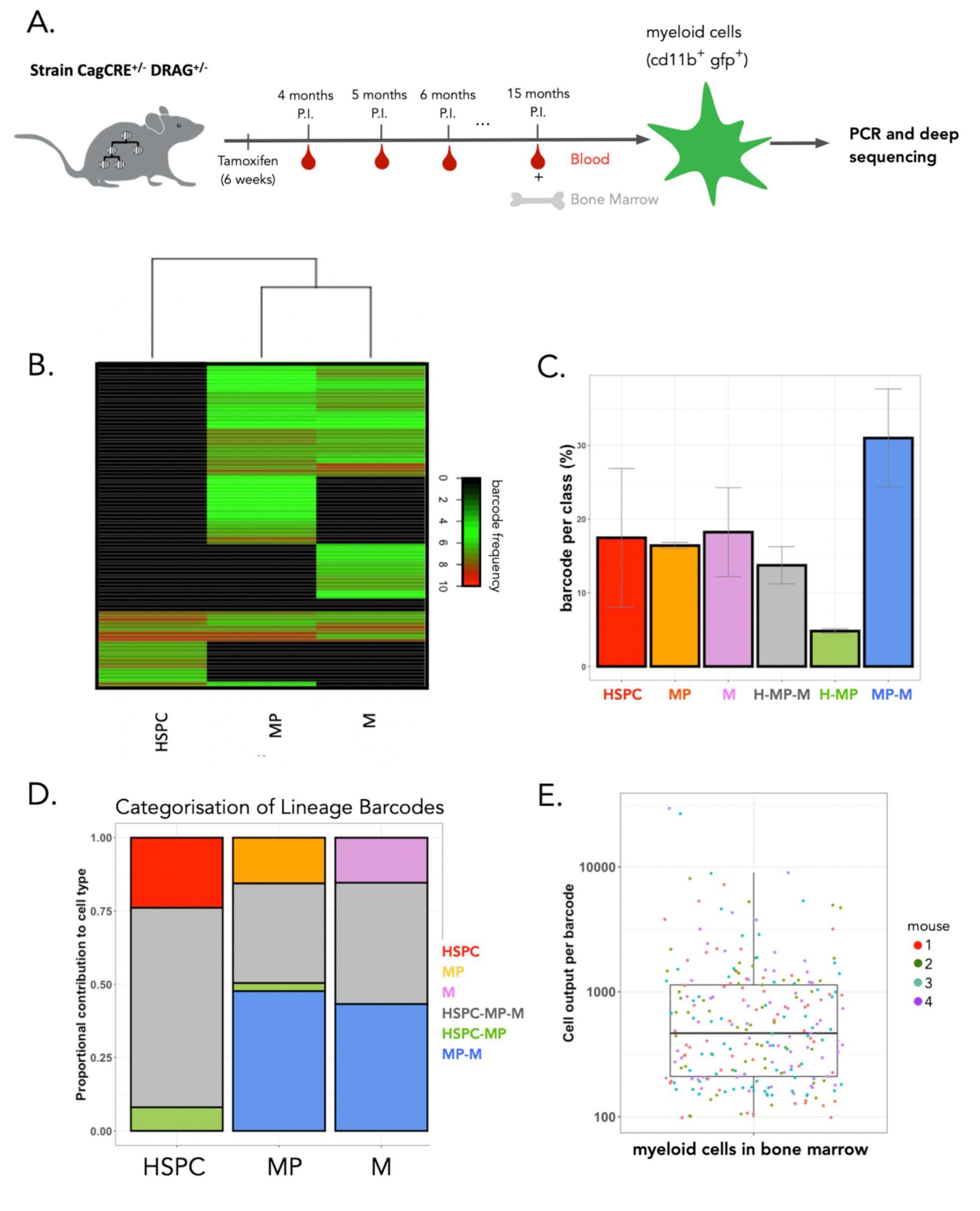
Barcode analysis reveals non-overlaping waves of myelopoiesis. (A) Recombination of the DRAG locus was induced in 8-14 week old mice. At sacrifice, 15 months post induction, myeloid cells (M: CDllb+ CD19·CD3·CDllc·) were sorted from bone marrow, HSPC (LSK: scal+ckit+) and myeloid progenitors (MP: myeloid progenitors, seal· ckit^+^) were sorted from the bone marrow (gating strategy in Fig. S4A and B). All samples were processed for barcode detection. B. Heatmap representation of the barcode output in bone marrow HPSC, MP, and myeloid cells, at month 15 post-induction. 97 barcodes with barcode generation probability Pgen < 10^−4^ were observed (pooled data of 3 mice). Normalized and hyperbolic arcsine transformed data were clustered by complete linkage using Euclidean distance. C. Barcodes with barcode generation probability Pgen < 10^−4^ were classified based on their presence or absence in HSPC, MP, or myeloid cells (M). The percentage of barcodes in each of the 6 possible classes is depicted. Error bars show the standard deviation between mice (n=3). D. Same as C but depicting the total contribution of all barcodes in a given class to the production of either HSPC, MP, or M. E. Number of myeloid cells produced per barcode for n=4 mice (251 barcodes), 15 months post-induction. Colors represent individual mice.

If not all HSPCs would be active at the same time, it may be expected that the numbers of clones in HSPCs could be higher than that detected in mature myeloid cells. To test for this prediction, total clone numbers were inferred from the observed diversity (chao2) numbers ^33^, taking into account labelling efficiency (chao2 analysis in Fig. S4D, S4E for other diversity estimates and Methods). Bone marrow myeloid cells were composed of at least 1,827 +/− 430 clones (mean and SD between mice), a value similar to published estimates ^19, 20^, while myeloid progenitors and HSPCs were composed of more clones, (2,646 +/− 1081 clones, and 3,040 +/− 953 clones respectively). This analysis suggests that not all clones present in the HSPC compartment are actively contributing to myelopoiesis, an observation that is compatible with a model of overlapping waves of myelopoiesis. Note that these estimates should be interpreted as the lower bound of total diversity, because of cell loss during extraction, limitation in the detection of small clones and the presence of recurrent barcodes. Together, these results suggest that steady state myelopoiesis is sustained by overlapping waves of HSPCs.

To explore whether individual long-term repopulating clones produce similar numbers of myeloid cells, we took advantage of our quantitative barcode detection system to determine the number of output cells per barcode clone. This analysis demonstrated that individual barcode-labeled long-term repopulating cells differ up to ~100 fold in their myeloid cell output (Fig. 3E), with clone sizes ranging from 100 to 10,000 cells. As a result of this, the majority of myeloid cells at an analyzed timepoint was produced by few barcoded long-term repopulating cells. This disparity in clone sizes may in part reflect temporal differences in clonal activity, but remains higher than the clonal diversity observed post-transplantation, in which ~ 100-fold fewer HSPCs are estimated to actively contribute to hematopoiesis ^34–37^. Overall, these data demonstrate that steady state myelopoiesis is the result of the unequal cellular output of a large number of clones, and are consistent with a model in which long-term repopulating cells maintains myelopoiesis through overlapping waves of cell production.

### Increase in the number of long-term repopulating cells contributing to myelopoiesis with ageing

To study the effect of ageing on the clonal composition and cell production of long-term repopulating cells, we sampled the blood of DRAG mice from month 4 until month 12 post-induction, as well as blood and bone marrow at month 15 post-induction (Fig. 3A). Due to limitations in the volume of blood that can be sampled at each timepoint, we observed a very low rate of barcode overlap between technical replicates in blood (9% +/− 8%, Fig. S6A) as compare to bone marrow samples (60% +/− 15%). Thus, within blood samples, only a small subset of all active clones are captured, precluding longitudinal analyses of individual clones, but still allowing one to follow changes in the number of contributing clones over time. Consistent with prior work ^1^, myeloid cell numbers in the blood of DRAG mice increased over time, and this increase was observed for both GFP^−^ and GFP^+^ (barcode labeled) cells (Fig. 4A). Likewise, an increase in total myeloid cells numbers between 6.5 and 19 month-old mice was observed in bone marrow (Fig. S7). Most bone marrow myeloid cells were neutrophils in both young (6.5 months) and aged mice (19 months) (young = 51.95 ± 0.7%; old = 64.6 ± 2.4%), and the frequency of neutrophils increased in old mice at the expense of macrophages and monocytes (p = 0.029) (Fig. S7). This result suggests an imbalance in the relative production rates of different innate immune cell subsets upon ageing.

**Figure 4:**
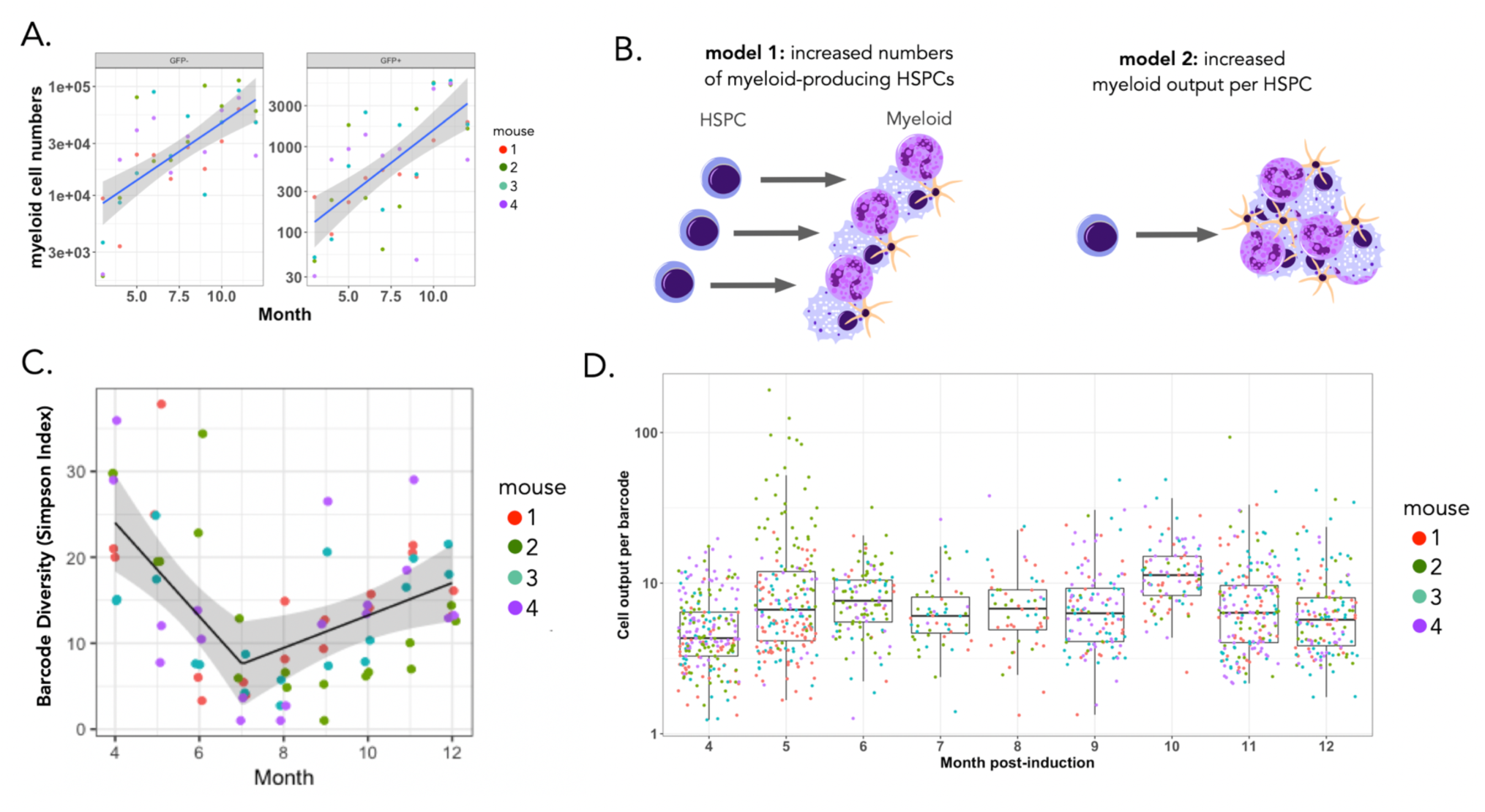
Increased numbers of long-term repopulating cells contribute to myelopoiesis with age. A. Absolute number of GFP^+^ and GFP^−^ myeloid cells (CD11b^+^) in blood between month 4 and 12. N=4 mice, black line depicts the mean, ribbon depicts the 95% confidence intervals for the true mean. B. Models for age-related increased myeloid cell production. An increase in myeloid production may happen through two non-mutually exclusive mechanisms: an increase in the number of myeloid-biased HSPCs (model 1), or an increase in the number of myeloid cells that are produced per individual HSPC (model 2). C. Diversity of barcodes in blood between month 4 and 12 using the Simpson index. Each sample was analyzed in duplicate. Black line represents the mean Simpson’s index estimate, obtained from the fitted gamma generalized linear mixed model with a break point; grey ribbon represents the 95% CI for the true mean. D. Number of myeloid cells produced per barcode (i.e. clone size) over time post-induction. Pooled data of four mice, with each color representing a different mouse, are depicted.

At the cellular level, the observed increase in myeloid cell production may occur by two non-exclusive mechanisms: an increase in the number of myeloid-producing long-term repopulating cells, or an increase in the number of myeloid cells that are produced per individual long-term repopulating cell (Fig. 4B). To distinguish between these scenarios, we analyzed the number of DRAG barcodes in the blood myeloid cell compartment over time. To this end, we computed the clonal diversity in sequential blood samples using Renyi entropy indexes (Fig. 4C) and modeled the change in diversity as a function of time using generalized mixed models, accounting for sampling (Suppl. Materials and methods). We found that a generalized linear mixed model with a break point showed the best fit to the experimental data (Fig. S6B). Applying this model to several diversity indices (Fig 4C, Fig. S6C, Table S9) showed a highly consistent trend over time. Specifically, in the first 7 months following DRAG barcoding, the number of clones contributing to the myeloid compartment decreased over time, consistent with the turn-over of shorter-lived cells that were labeled using the ubiquitous CagCre driver (Fig. 2A). Strikingly, after this time point, the number of barcodes contributing to myelopoiesis increased linearly (Fig. 4C), indicating that the number of long-term repopulating cells contributing to myelopoiesis was increasing over time. This increase could not be explained by the slight preferential labeling of MPP3, as MPP3 numbers did not increase with age (Fig. 5D). Furthermore, increased barcode diversity was also not explained by a delayed recombination of DRAG barcode V regions, as the majority of barcodes was associated with a single unique V region, and as the frequency of unique V regions remained constant over time (Fig. S6D). Strikingly, the number of myeloid cells produced per long-term repopulating clone did not change over time (Fig. 4D). Collectively, these results reveal that the increase in myeloid production upon aging is due to an increased number of myeloid producing long-term repopulating cells, rather than an increased clonal output of individual long-term repopulating cells.

**Figure 5:**
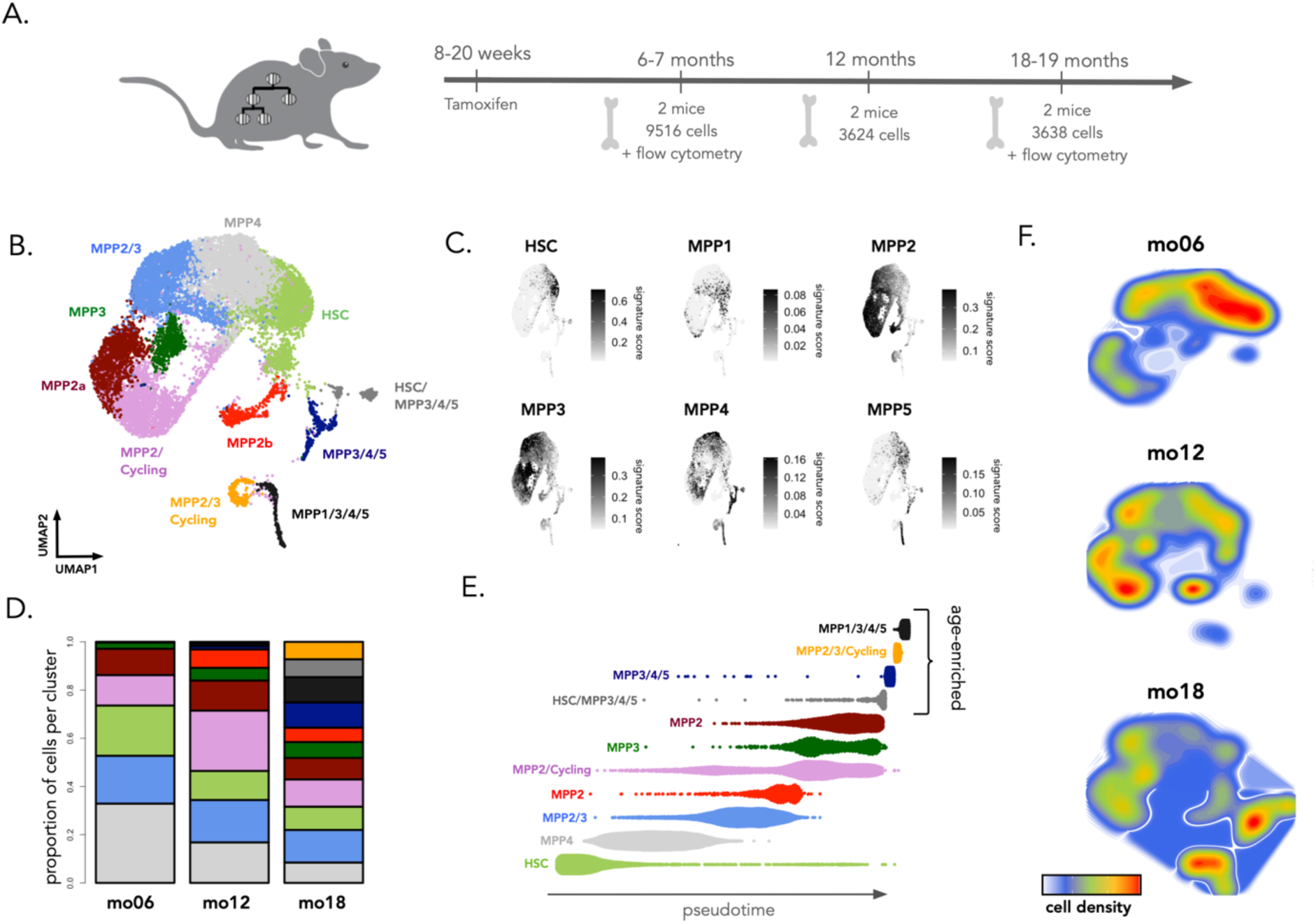
Age-related changes in cellular composition of the HSPC compartment. (A) Experimental timeline for profiling HSPCs across adulthood. Mice were given tamoxifen at 8-20 weeks of age to induce barcode recombination. At subsequent timepoints HSPCs were purified from the bone marrow of induced mice and processed for scRNAseq or flow cytometry analysis. For each timepoint we give the number of mice processed as well as the number of cells recovered for scRNAseq profiling. (B) UMAP embedding of the scRNAseq data. Unsupervised clustering was used to discretize the data into colored subgroups and cluster annotation was performed by overlaying published gene signatures and markers ^27, 28, 59^. (C) Overlaying published gene signatures ^28, 59^ onto the UMAP embedding of the data. For each cell the gene signature score was calculated as the mean expression across all genes in the signature (after background correction). (D) The proportional abundance of cells amongst clusters at 6-7 months, 12 months and 18-19 months old. (E) Pseudotime projection of the data with cells organized into clusters as in C. Pseudotime inference was performed using a diffusion map based approach as implemented in the R-package destiny ^42^. (F) Density plot showing the proportional abundance of cells within the UMAP embedding as a function of age.

### Age-related transcriptomic changes in HSPCs

Our *in situ* lineage tracing analyses show that ageing leads to an increase in the frequency of long term repopulating clones that actively contribute to myelopoiesis. To understand the cellular and molecular processes that give rise to this phenomenon, we performed single cell transcriptomic (scRNA-seq) profiling of Sca1^+^ cKit^+^ GFP^+^ HSPCs purified from mice aged 6,5, 12, and 19 months old (n = 6 mice) (Fig. 5A). Flow cytometry analysis of GFP labelling in HSPCs from aged (19 months) mice showed no significant differences in GFP^+/−^ proportions amongst the HSC and MPP1-5 subsets, confirming that the DRAG barcoding system does not preferentially label a specific HSPC subset, even in aged mice (Figure S12B). Following scRNAseq data quality control and pre-processing, we recovered the transcriptome for 16,778 cells with a median of 2,845 genes detected per cell. After data integration and non-linear dimensionality reduction (UMAP), we performed unsupervised clustering of the data, and annotated the 11 resultant clusters using published gene signatures and markers ^27, 28, 32^ (Figure 5B-C, Figure S8A-B, S9A, Table S13). Consistent with reports indicating that HSPCs do not form discrete cell subsets ^38–41^, we observed that many clusters co-expressed signatures of the MPP1-5 subtypes (Figure 5C, S8B). In cases where clusters could not be assigned to a single HSPC subset, we named the cluster according to the different combinations of HSC and MPP signatures that they expressed ^27^. Using this reference embedding and supervised annotation of the data, we observed an accumulation of transcriptomic changes throughout adulthood (Fig. 5D-F), with several clusters enriched in either young or in aged mice (Fig. 5D, 5F). Specifically, these analyses suggest that LT-HSCs are less frequent in aged mice (Fig. 5D), whereas 4 clusters were only found in aged mice (Fig. 5D, 5F). A common feature of these ageing-associated clusters was the co-expression of the MPP3, MPP4 and MPP5 gene signatures (Fig. 5C-D) and expression of genes associated with mature myeloid cells (CD74, Ngp, Cll5, Fig. S9A). Pseudotemporal ordering of the data using a diffusion map approach ^42^ predicts that HSPCs from aged-associated clusters represent a more differentiated cell state as compared to HSPCs from young mice (Fig. 5E). In addition, using a supervised annotation approach in which we mapped our cell clusters onto an independent reference dataset ^43^ (44,802 c-kit^+^ and c-kit^+^ sca1^+^ cells), we observed an age-related overall reduction in the number of cells that mapped to LT-HSCs, while cells from age-associated clusters increasingly mapped to lineage restricted progenitors including, GMP, MEP and CLP (Fig. S9B-C). Together these data show that ageing leads to the reduced expression of genes associated with quiescent LT-HSCs, and the emergence of MPP-like cell states with features of enhanced differentiation.

We then further characterized the transcriptomic changes accumulating with age in particular in the age-associated MPP-like cell states. To understand if ageing leads to a change in cell cycle rates, cells were classified into G1/G2+M/S phases of the cell cycle based on their gene expression patterns ^44^ (Fig. 6A, Fig. S9E). We observed age-associated increases in the proportion of cells in G2/M and S phases of the cell cycle across multiple clusters, including the age associated MPP3/4/5, MPP1/3/4/5, HSC/MPP3/4/5 and the MPP2/3/Cycling clusters (Fig. 6A). Differential gene expression analysis between HSPCs of different ages showed an overall increased expression of myeloid associated genes *S100a8, S100a9, Elane, Mpo and Fcer1g* with age and decreased expression of genes associated with LT-HSCs including *Procr, Ltb* and *Tcf15* ^45, 46^ (Fig. 6B, Table S14). Differential expression and gene-set enrichment analyses also showed that aged HSPCs have increased expression of genes related to inflammation, cytokine stimulation, cycling, and DNA damage, in line with prior data ^4, 5, 12, 13^, as well as transcriptomic changes related to the regulation of protein ubiquitination and the electron transport chain (Fig. 6C, Fig. S10,Table S14-15). To assess which specific compartments were most affected by ageing we aggregated genes upregulated in aged HSPCs (from 19 month old mice) into an aged HSPC signature and assessed its expression across all clusters (Figure 6D). This analysis showed that much of the transcriptomic differences between young and aged HSPCs occurred within age-associated MPP compartments and MPP3, rather than HSCs or other MPPs (Figure 6D). Collectively, these scRNAseq analyses suggest that ageing leads to a decreased number of LT-HSCs and the emergence of age-associated MPP3/4/5-like cell states. Transcriptomically, these aged HSPCs display features of increased cycling, stress and metabolic gene expression. Together with our functional DRAG barcoding analyses, our data suggests that while HSPCs in the native bone marrow accumulate transcriptomic changes with ageing, their myeloid production rates remain consistent over time, suggesting that these changes are not impacting their ability to produce cells.

**Figure 6:**
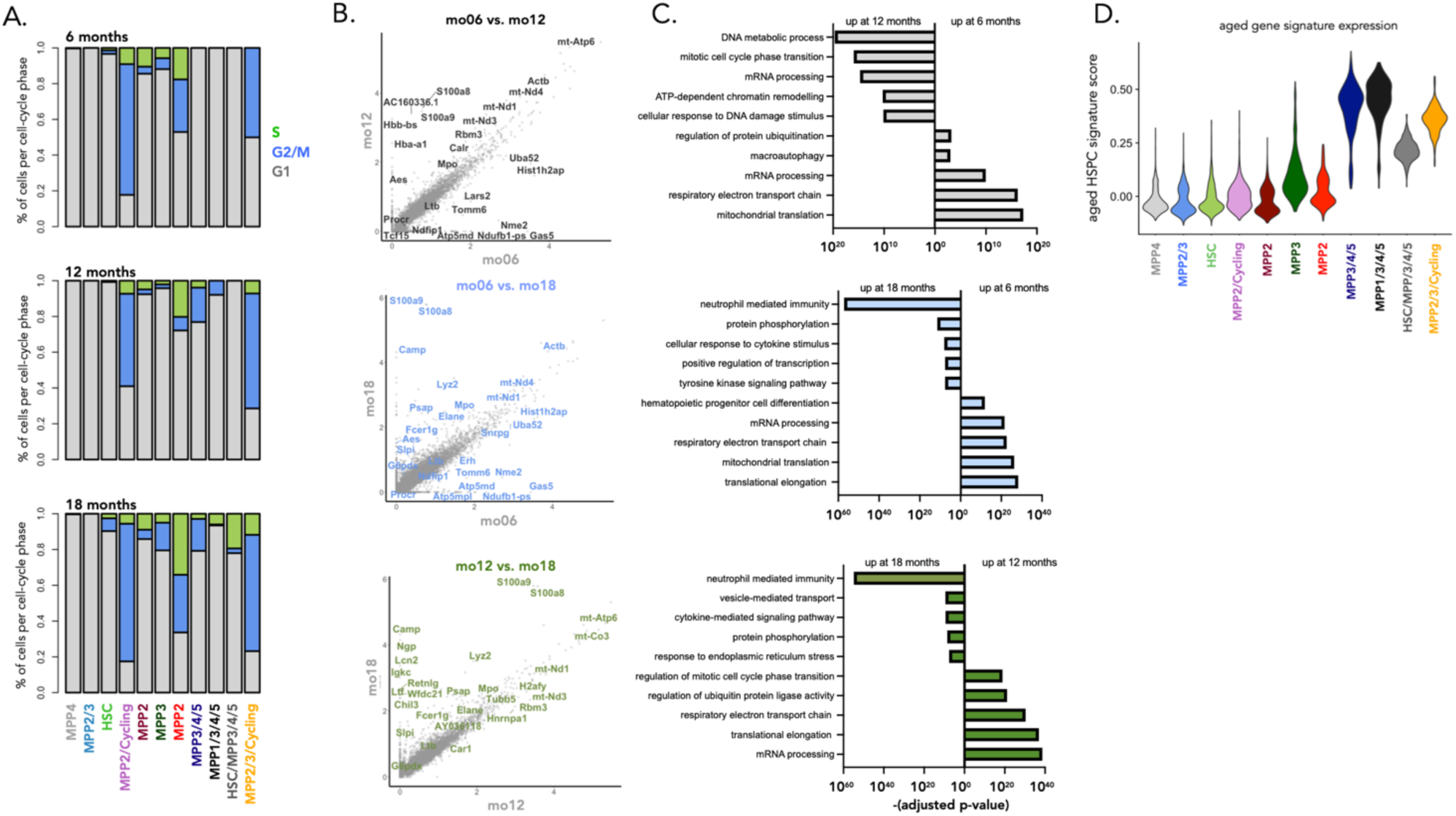
Transcriptomic differences between young and aged HSPCs. A. Proportion of cells per cell cycle phase, per cluster and age. Cells were classified into G1/G2+M/S phases of the cell cycle using the classifier approach developed by ^44^. B. Differentially expressed genes between HSPCs from mice aged 6.5, 12 and 19 months. Differential expression analysis was performed using a logistic regression test as implemented in the Seurat R package. Bonferoni correction was applied to correct for multiple testing C. Pathways enriched in HSPCs at different ages. Pathway analysis was performed using the enrichR R package using a variation of Fisher’s exact test, which also considers the size of each gene set when assessing the statistical significance of a gene set ^60^. D. Expression of the aged HSPC gene signature across all cell clusters. The gene signature was obtained by differential expression analysis for all HSPCs between young (6.5 months) and aged (19 months) mice. Genes that are upregulated in aged HSPCs are aggregated into the aged HSPC signature. For each cell, the gene signature score was calculated as the background corrected mean expression across all genes in the signature.

## Discussion

In this work, we present a new *in situ* DRAG barcoding system that allows for efficient, neutral, stable, and diverse labeling of individual hematopoietic cells. In addition, the quantitative detection of resulting barcodes in downstream cells makes it possible to enumerate clonal output. Relative to other barcoding strategies (Table S12), the DRAG system offers several advantages such as a straightforward PCR and sequencing strategy (through the use of UMI and Single Read 65bp illumina sequencing), and a quantitative framework to filter and analyze barcoding data with high resolution. We do note that the utility of the approach in the lymphoid system is restricted because of the expected cassette recombination during B and T cell development. The DRAG barcoding system should be of value to examine aspects of tissue generation in other cell systems, such as brain and mammary gland, in which efficient labeling with limited background is observed (Fig. S11).

Using the DRAG barcoding system, we observe that HSPCs are highly heterogeneous in the time and extent they contribute to myelopoiesis, with relatively few barcodes shared across the entire myeloid developmental trajectory, and clone sizes varying by several orders of magnitude. Clone size variability was not fully explained by differences in when clones were active as large clone size variations were observed in clones that were active at the same time (clones that were found across the entire myeloid developmental trajectory - HSPC, myeloid progenitors and mature myeloid) (Fig. S4G). Our results extend previous findings on the existence of differentiation-inactive^47^ or childless^48^ HSPC from other barcoding studies in the native niche by showing quantitative heterogeneity in cell production by HSPC. Together, these data support a model in which myelopoiesis is sustained by overlapping waves of HSPC activation, with large variations in the cellular outputs of each differentiation-active HSPC clone.

Ageing of the immune system is associated with dramatic changes in the distribution and functional properties of immune cells. Broadly, there is a skewing in favour of innate vs adaptive immunity, leaving older individuals increasingly susceptible to infection and chronic tissue inflammation. A lack of tools capable of measuring the output of individual HSPCs in the native bone marrow microenvironment has complicated the research literature around this topic ^18^, with many associations being drawn between phenotypic changes in native hematopoiesis as observed by scRNAseq^14, 49^ and functional changes in cellular output as observed in post-transplantation hematopoiesis ^6–10^. In transplantation assays, aged HSPCs display a skewed output towards the myeloid and platelet lineages^3, 6, 7, 15, 16^, have a lower rate of self-renewal^3, 8, 15^ and have a decreased cell production capacity^1, 15^ relative to young HSPCs. Coupling of these transplantation-based functional measurements with gene expression patterns associated with stress and inflammation in native hematopoiesis have led to a model in which aged HSPC exhaustion is a hallmark of an ageing immune system^17^. However, our results on native hematopoiesis upon ageing do not fully support this model and are consistent with studies in young mice showing that native hematopoiesis differs from post-transplantation hematopoiesis ^18^. Specifically, using longitudinal monitoring of the diversity of endogenous DRAG barcodes in blood we found that the increased myeloid production occurs through an increase in the number of long-term repopulating clones, rather than through an increased number of myeloid cells produced per clone. We therefore conclude that while aged HSPCs do exhibit transcriptomic signs of cell stress, inflammation and changes in global gene expression state, these cells are still able to functionally produce the same amount of myeloid cells, contradicting the current view that HSPC in their native niche are dysfunctional in their cell-production capacity.

Not all HSCs are differentiation-active at the same time during adulthood, as shown by our findings in this study and from previous reports^47, 50^, raising the question on whether the well-documented increase in phenotypic HSC numbers with ageing corresponds to an increase in differentiation-active or inactive HSCs. Transplantation studies suggest that part of this increase corresponds to an increase in the number of differentiation-active HSCs. However transplantation assays do not inform us on whether the HSPCs that accumulate with age are actively contributing to regeneration because of the perturbation generated by transplantation. Here, we show that the number of HSPC clones actively contributing to myelopoiesis increases with age in the native bone marrow. Importantly, this increased number of differentiation-active HSPCs could favor the occurrence of genetic mutations associated with clonal hematopoiesis^51–53^, increasing the risk of hematological malignancies associated with age.

The finding that differentiation-active HSPCs increase in number with age is explained at the cellular level by a decreased number of LT-HSCs and the increase in frequency of cycling MPPs as observed by scRNAseq profiling, and suggest that LT-HSCs in aged individuals exit quiescence and contribute to hematopoietic flux at a faster rate than in younger individuals. The increased entry into cycling and differentiation of HSC could potentially be caused by repeated exposures to inflammation over the course of adulthood, and the occurrence of age-associated MPPs with signs of cell stress and inflammation forms indirect evidence for such a model. Furthermore, evidence that inflammation pushes LT-HSCs to differentiate ^54–56^ and that ageing induces proliferative JAK/STAT signaling in HSPCs ^57^ are in line with this proposed mechanism. Of note, the changes in HSPC composition from scRNAseq were not consistent with changes in population dynamics when defined using surface markers (Fig. S12), suggesting that further work is needed to develop a unified definition and nomenclature for HSPCs that is consistent throughout adult and aged hematopoiesis. As the bone marrow niche has also been shown to change upon ageing ^58^, extrinsic factors may also contribute to the increased number of active long-term repopulating clones in aged mice.

In summary, our study highlights the utility of quantitative in situ barcoding methods, and suggests that greater caution should be exercised when extrapolating results from transplantation assays to native hematopoiesis. Our data does not support a model in which aged HSPCs are dysfunctional in their cell production capacity.

## Acknowledgements

We would like to thank R. Bin Ali for blastocyst injections, Dr. R. Gerstein, Dr. K Vanura, and Dr. S. Gilfillan for sharing reagents, and M. Hoekstra for sharing drawings. We thank past and present members of the Schumacher lab, in particular Dr. S. Naik, Dr. C. Gerlach, and Dr. J. Rohr, for valuable discussions. We thank Dr. K. Duffy, Dr. R. de Boer, Dr. L. Riboli-Sasco, Dr. P. Krimpenfort, Dr. J Jonkers and the Perié team for helpful discussions. We thank the Curie flow cytometry, next-generation sequencing, and animal facility from both NKI and Institut Curie.

## Data and materials

availability: all data and scripts are available on the gitlab of the Perié team.

## Contribution

LP, JU and JC designed and performed experiments. LP supervised the study, analyzed data and created figures with help from JU, JC and JB. CC, AMW, ET contributed to experiments, TS conceived the technological approach, JVH, HJ, JU, and TS designed the DRAG recombination substrate and mouse. LK and JU isolated MEF clones. CM performed and analyzed the mammary gland experiment, LP, CC and JM performed the brain experiment. SF and JF supervised the mammary gland and brain experiment respectively.JB analyzed MEF data and developed the filtering pipeline with input from LP, JU and TS. AV designed the preprocessing pipeline. JC designed and performed the scRNAseq analysis and brain data. YE, AMW, and TM designed the probability generation model. RAM designed the generalized linear mixed models. LP and TS wrote the main text of the manuscript with feedback from all authors.

## List of Supplementary Materials

Materials and Methods

Table S1 – S15

Fig S1 – S10

Sup References

## Materials and Methods

### DRAG construct

A GFP-based VJ recombination substrate ^1^ (kindly provided by Dr. R. Gerstein, University of Massachusetts Medical School, USA) was inserted into the plasmid pSBEX3IB. To assemble the DRAG substrate, a “12 RSS-J segment” fragment was generated by PCR and cloned into pBluescript using *Sac*I and *Kpn*I. An eGFP encoding gene fragment was ligated 3’ of the J-segment using *Nco*I and *Sac*II. Next, a “V-segment-12 RSS” fragment was generated by PCR and inserted 5’ of the J-segment using *Sac*I and *Eco*RI. Then, a “23 RSS-D segment-23 RSS-bovine growth hormone (BGH) polyA signal” fragment was generated by PCR and cloned in between the V- and J-segment using *Bgl*II and *Eco*RI. Spacer sequences of the two D-segment 23-RSSs were varied to prevent hairpin formation. D-segment sequence is the naturally occurring IgH DSP2.4, in which the naturally occurring ATG sequence was mutated into ATC, to prevent premature translational initiation. The complete “V-segment-12 RSS-23 RSS-D-segment-23 RSS-BGH polyA-12 RSS-J segment-GFP” fragment was cloned into a Rosa26 targeting vector containing a CMV enhancer and chicken beta-actin promoter using *Asc*I. Finally, the “loxP-TdT-E2A-RAG2-T2A-RAG1-loxP” cassette was inserted in antisense orientation 3’ of the V-segment, using *Pml*I and *Mfe*I, to give rise to the DRAG targeting construct as depicted in Fig. 1A.

### Quantifying the sensitivity and specificity of the DRAG barcoding system

Taking advantage of the VDJ recombination system that produces a high degree of genetic diversity in the lymphoid lineages, we designed a DNA cassette, termed DRAG (Diversity through RAG), with the aim to allow endogenous barcoding of all cellular lineages in an organism in a temporally controlled manner (Fig. 1A). The DRAG system has been designed such that upon CRE induction, a segment between two loxP sites is inverted, leading to the expression of both the RAG1 and 2 enzymes and Terminal deoxynucleotidyl transferase (TdT). Upon such expression, recognition of recombination signal sequences (RSSs) within the DRAG cassette by the RAG1/2 complex leads to recombination of the synthetic V-, D- and J-segments, with diversity being generated both by nucleotide deletion and TdT-mediated N-addition. Notably, as the RAG/TdT cassette and RSSs are spliced out during this recombination step, further recombination of the DRAG locus is prevented, and any generated VDJ sequence is thus stable over time. Finally, recombination of the DRAG locus results in the removal of a BGH polyA site that precludes GFP expression in the native DRAG configuration, allowing one to identify barcode^+^ cells by flow cytometry or imaging. Note that the GFP expression is driven by the CAGGS promoter upfront of the barcode to link its expression with the presence of a barcode.

To understand DRAG recombination patterns, we isolated MEF cells from DRAG mice, induced DRAG recombination with tamoxifen, and derived MEF clones (n=24) that carried a single DRAG barcode by limited dilution. In line with expectations, recombined DRAG loci were characterized by insertions and deletions between the VDJ segments (Table S1), and only 2 over 24 barcodes were shared. In addition, consistent with the DRAG design, longitudinal analysis of barcode sequences showed that identified barcodes were stable over time, as assessed by Sanger and deep sequencing, for 7 out of 7 MEF clones tested. The barcodes that are generated upon DRAG recombination are random, precluding the use of barcode reference lists to distinguish true recombination events from amplification and sequencing errors. To address this issue, we developed a DRAG barcode processing platform that incorporates unique molecular identifiers (UMI) during PCR amplification (Fig. S1B Sup Method) and a strict filtering pipeline of sequencing data. Application of this strategy to defined mixtures of 7 MEF clones that each contain a different barcode demonstrated that this platform allows the robust detection and quantification of barcodes (Fig. 1D). Specifically, clones equal or greater than 10 cells were efficiently and quantitatively identified in samples with few cells (97 cells), and clones equal or greater than 100 cells were efficiently and quantitatively identified in samples with larger cell numbers (25,000 cells), the latter corresponding to detection of clones that make up as little as 0.4% of the total cell population. The number of false positive events was 1 or, in most cases, 0. Importantly, across the range of detection, experimentally observed frequencies were highly correlated to input frequencies (Fig. 1D), demonstrating the ability of DRAG to allow quantitative analyses of cellular output.

To explore the genetic diversity obtained by in vivo DRAG barcoding, we next analyzed barcode diversity in myeloid cells 15 months post induction of DRAG recombination. Deep sequencing analysis of resulting barcode sequences revealed that DRAG barcodes characterized by up to 15 nucleotide insertions and 32 nucleotide deletions were formed (Fig. S2A) and that the D-segment was inverted in 12% of cases.

In any system that creates semi-random codes for lineage tracing, the information value of such codes depends on the probability of their occurrence, as barcodes with a lower likelihood of generation form more reliable single cell identifiers ^2^. Applying a mathematical model ^3^ to infer the probability of generation of each barcode (P_gen_) from the data (Suppl. methods and Table S4–9), we observed a distribution of P_gen_ that spanned over at least ten orders of magnitude (Fig S2B). To experimentally test the values of the P_gen_ estimates, we analyzed the occurrence of shared and unique barcodes across a cohort of four tamoxifen-induced DRAG mice. Importantly, barcodes with a low P_gen_ estimate were in almost all cases unique to each mouse (Fig. S2B). In contrast, barcodes with a high Pgen estimate were generally shared between mice (Fig. S2B) and also had a higher average read frequency (Fig. S2C), consistent with their independent occurrence in multiple progenitor cells. On the basis of these results, we selected a P_gen_ estimate of <10^−4^ for further data analysis (Fig. S2D-E). At this probability cut-off, 61% of barcode sequences were retained, and 92% of the retained barcodes were unique to an individual mouse. In addition, using this analysis strategy, hundreds of barcodes that could be used for analysis were retained for each mouse. Together, these data show that the DRAG mouse model allows the in vivo creation of a large barcode diversity without a requirement for cell transplantation, and that the prevalence of these barcodes in downstream progeny can be detected in a quantitative manner.

#### Mice

All animal breeding and experiments were performed in accordance with national guidelines and were approved by the Experimental Animal Committee of the NKI (DEC 09036) or Institut Curie (#16854-2018092412148925-v1).

#### DRAG mice generation

The DRAG targeting construct was linearized using *Pvu*I and electroporated into IB10 E14 129/ola ES cells. Stable transfectants were selected with puromycin and resistant clones were picked and expanded. Correct integration was determined by Southern blotting, using a probe directed against the 5’ Rosa26 homology arm. Two independent ES cell clones were injected into C57Bl/6 blastocysts to generate 28 transgene-positive chimeric mice and establish two independent DRAG transgenic lines (DRAG1 and DRAG2). GFP expression in peripheral blood B and T cells was screened using anti-CD19-PE (BD, clone 1D3, dilution 1/100), anti-CD3e-PerCP-Cy5.5 (eBioscience, clone 145-2C11, 1/100) and anti-CD11b-APC (BD, clone M1/70, 1/100), and the DRAG1 line was selected. DRAG1 mice were crossed with B6.Cg-Tg(CAG-cre/Esr1*)5Amc/J (CAGGCre-ER^TM^) to obtain heterozygous mice for experimental use.

#### Tamoxifen induction

Six to 20 week old male mice received 7 mg Tamoxifen/ 40 gr bodyweight each day for 3 consecutive days by intraperitoneal injection. Tamoxifen (T5648-1G, Sigma) was dissolved 10% EtOH and 90% sunflower oil (Sigma).

#### Blood sampling

100-200µl tail vein blood samples were obtained at the indicated time points. At sacrifice, a larger blood volume was obtained by heart puncture.

#### Single barcoded MEF clones

Embryos from DRAG1 homozygote CagCreERT2 homozygote mice were harvested at day 14.5 and PCR genotyped. Extracted MEFs were immortalized by transduction with p53shRNA (in pRetro-super backbone, kindly provided by M. v. Lohuizen) and puromycin selected. Recombination of the DRAG transgene was induced in vitro using 5 µM 4-OH-Tamoxifen. Following induction, single GFP+ cells were grown by limiting dilution. Clones were individually Sanger sequenced to identify DRAG barcodes. To confirm barcode identities, clones were also deep-sequenced using the same PCR pipeline (including capture) as used for other DRAG samples (see below). To create MEF mixes, 7 clones, each harboring a different barcode, were mixed in a ratio of 64:32:16:8:4:2:1. Resulting mixtures were used to prepare pools of 50,000, 12,500, 3,125, 781 and 195 cells, and samples were further processed as described for DRAG samples.

### Hematopoietic cell isolation and sorting for DRAG analysis

100-200 µl blood samples were directly harvested in 800 µl Erylysis buffer (80.2 g NH_4_Cl, 8.4g NaHCO_3_, 3.7 g disodium EDTA in 1 L H_2_O, pH 7.4), incubated on ice, diluted with 5 ml Erylysis buffer, washed with medium RPMI, resuspended in 0.5 ml 10% RPMI medium and put on ice overnight. Subsequently, blood cells were stained in 50 µl 2% FCS RPMI medium with antibodies against CD11C (APC, clone HC3, BD biosciences, dilution 1/100), CD11b (PercPCy5.5 or Pacific Blue, clone M1/70, ebioscience, 1/100), CD19 (APC-Cy7, clone 1D3, BD Pharmingen, 1/100) and CD3 (percp 5.5, clone 145-2C11 or PE, clone eBio500A2, eBioscience, 1/100). At sacrifice, BM was harvested from femurs, tibias and ilia and cells were enriched using anti-CD117 magnetic beads (Miltenyi). The c-kit^+^ fraction was stained with antibodies against CD117 (c-kit APC, clone 2B8, Biolegend, 1/100), Sca-1 (Pacific Blue, clone D7, eBioscience, 1/200), CD135 (Flt3 PE, clone A2F10, ebiosciences, 1/100) and CD150 (Slam Pecy7, clone TC15-12F12.2, biolegend, 1/100). The c-kit^−^ fraction was stained with antibodies against CD11C (APC, clone HC3, BD biosciences, 1/100), CD11b (PercPCy5.5 or Pacific Blue, clone M1/70, ebioscience, 100), CD19 (APC-Cy7, clone 1D3, BD Pharmingen, 1/100). When two colors are indicated for a given antibodies, they were used for the same cohort but at different time points for all mice. Cells were sorted on a FACSAria^TM^ (BD Biosciences) in FCS coated eppendorf tubes, according to the gating strategy presented in Fig. S4C and D.

#### Barcode PCR and deep sequencing

**Lysis:** Sorted cells were lysed in 40 µl DirectPCR Lysis Reagent (Cell) from Viagen Biotech with 0.4 mg Prot K, and incubated for at least 1 h at 55°C, followed by a 30 min heat inactivation at 85°C and 5 min at 94°C. Samples were stored at −20°C. **Shearing genomic DNA (gDNA):** samples were complemented up to 130 µl with 10 mM Tris, and shearing was performed on a ME220 Focused-ultrasonicator (Covaris) in 130 µl reaction tubes under the following conditions: time: 20 sec; peakpower:70; duty% 20; cycles/burst:1000. **Capture:** sheared gDNA of each sample was split in two duplicates and samples were incubated o/n at 65°C after mixing with an equal volume hybridization buffer (1 ml composition: 667 µl 20x SSPE (Gibco); 267 µl 50x Denhardt’s solution (Sigma-Aldrich); 13.3 µl 20% SDS (Sigma-Aldrich); 26.7 µl 0.5M EDTA (Sigma-Aldrich); 26.7 µl nuclease free water (Ambion), together with capture oligos (50 fmol each). The next day, 5 µl streptavidin beads (Dynabeads^tm^ MyOne^tm^ streptavidin T1) were washed twice with 100 µl 2X B&W Buffer (2M NaCl in TE buffer) in pre-rinsed (with 400 µl 10 mM Tris solution) low retention microtubes (axygen). The Biotinylated gDNA was mixed with the beads in an equal volume 2X B&W Buffer and samples were incubated for 30 min at RT, mixed every 10 min. Beads were washed subsequently in:-500 µl 1X B&W Buffer(2X B&W diluted with TE buffer);-200 µl ½x b&w Buffer(2X B&W diluted with TRIS buffer);-75 µl ¼x b&w Buffer(2X B&W diluted with TRIS buffer); twice in 75 µl 10 mM TRIS buffer. **Preamp PCR:** Beads were resuspended in 200 µl PCR mix (20 µl 5x phusion HF buffer (NEB); 1 µl Phusion DNA Polymerase (NEB); 2 µl 10 mM dNTPs; 0.5 µl 100 uM preamp forw. oligo; 0.5 µl 100 uM preamp rev. oligo; 76 µl PCR grade water) and split in two replicates. PCR program: 2 min. at 98°C; N* cycles of 10 sec at 98°C, 20 sec at 60°C, 25 sec at 72°C; 5 min. at 72°C; 4°C forever * The number of cycles (N) is adjusted according to number of barcoded cells in the sample, such that every sample has the same number of molecules at the start of the tagging PCR (See Table S3 for cycle numbers used). **Tagging PCR:** 2 µl preramp PCR product was mixed with 48 µl tagging PCR mix (10 µl 5x phusion HF buffer (NEB); 1 µl Phusion Hot Start II DNA Polymerase (2U/ µl) (NEB); 1 µl 10 mM dNTPs; 0.25 µl 1 µM M1 tag forward oligo; 35.75 µl PCR grade water). PCR program: 1 min. at 98°C; 2 cycles of 10 sec at 98°C, 2 min. at 57°C, 1 min. at 72°C; 20°C forever). To digest remaining M1 tag forward oligo: 3 µl 20U/ µl Exonuclease I (NEB) was added and samples were incubated for 1 h at 37°C ExoI denaturation was performed for 5 min at 98°C, and samples were cooled down to 20°C. 0.5 µl 100 µM Illumina forward seq oligo and M1rev oligo (both with 2 Phosphorothioate bonds at the 3’end to prevent breakdown because of residual ExoI activity) were added, followed by PCR program: 1 min. at 98°C; 30 cycles of 10 sec at 98°C, 20 sec at 67°C, 25 sec at 72°C; 5 min. at 72°C; 4°C forever. **Sample index PCR:** 2 µl tagging PCR product was mixed with 18 µl PCR mix (4 µl 5x phusion HF buffer (NEB); 0.4 µl Phusion DNA Polymerase (NEB); 0.4 µl 10 mM dNTPs; 0.1 µl 100 uM P5 forw. oligo; 4 µl 2.5 uM P7 index rev. oligo; 9.1 µl PCR grade water). PCR program: 30 sec at 98°C; 15 cycles of 10 sec at 98°C, 20 sec at 67°C, 25 sec at 72°C; 5 min. at 72°C; 4°C forever). **Deepseq analysis:** 10 µl from each index PCR product was taken and pooled (140 samples per deepseq run), cleaned and concentrated 20x using the Monarch PCR & DNA Cleanup Kit (5 µg) (NEB). Further cleanup was done with the E-gel imager system (Invitrogen) and the extracted PCR product mix was concentrated to its original volume by means of speedvac concentration and analyzed on a 2100 Bioanalyser instrument (Agilent). Pooled samples were deep-sequenced on a HiSeq 2500 System (Illumina) in SR100bp Rapid run mode.

### Oligo sequences

#### Mef clones 2^nd^ round PCR for Sanger sequencing

M1seqv2for: ACACTCTTTCCCTACACGACGCTCTTCCGATCTNNNNCTCGAGGTCATCGAAGTA TCAAG

SM2index v2: GTGACTGGAGTTCAGACGTGTGCTCTTCCGATC CGTCCAGCTCGACCAGGAT

#### Capture

Biotinylated capture forward:

5’-BiotinTeg-CCGCTAGCGGCCAGGGCGGCCGGAGAATTGTAATACGACTCACTATAGGGAGACGCGTGTTACCTCCTCGAGGTCATCGAAGTATCAAG

Biotinylated capture:

5’-BiotinTeg-CTATAGCGGCCGCCTAGGCCGCTCTTCAACTACC TTGTACAGCTCGTCCATGCCGAGAGTGATCCCGGCGGCGGTCACGAACTCCAGCA GGACCATGTGA

#### Tagging PCR

Preamp forward: ACTCACTATAGGGAGACGCGTGTTACC Preamp reverse: GACACGCTGAACTTGTGGCCGTTTA

M1 tag forward: ACACTCTTTCCCTACACGACGCTCTTCCGATCNNNNNNNNNNNNCCTCGAGGTCA TCGAAGTATCAAG

Illumina forward seq (Read 1): ACACTCTTTCCCTACACGACGCTCTTCCGA*T*C (*= Phosphorothioate bond)

M1 rev Read2: AGTTCAGACGTGTGCTCTTCCGATC CAGCTCGACCAGGATG*G*G P5 forward : AATGATACGGCGACCACCGAGATCTACACTCTTTCCCTACACGACGCTCTTCCGA TC

P7 index rev: CAAGCAGAAGACGGCATACGAGAT**XXXXXXX**GTGACTGGAGTTCAGACGTGTGC TCTTCCGATC

Complete list of the i7 indexes in table S11.

All oligos were ordered at IDT with HPLC purified grade.

### Hematopoietic and myeloid cell composition analysis by cytometry

BM was harvested from femurs, tibias and ilia and cells were enriched using anti-CD117 magnetic beads (Miltenyi). The c-kit^+^ fraction was stained with antibodies against CD117 (c-kit APC, clone 2B8, Biolegend, dilution 1/100), Sca-1 (APC-Cy7, clone D7, Biolegend, 1/100), CD135 (Flt3 PE-Cy5, clone A2F10, Life technologies, 1/50), CD150 (Slam Pecy7, clone TC15-12F12.2, Biolegend, 1/100), CD48 (Pacific Blue, HM48-1, Biolegend, 1/100), CD16/32 (PercPCy5.5, clone M1/70, ebioscience, 1/100), CD34 (Alexa 700, 1/100) and a lineage cocktail (PE, CD3ε clone 145-2C11; Ly-6G/Ly-6C clone RB6-8C5; CD11b clone M1/70; CD45R/B220 clone RA3-6B2; TER-119, Biolegend, 1/200). The c-kit^−^ fraction was stained with antibodies against CD11b (PercPCy5.5, clone M1/70, ebioscience, 1/100), Ly6C (APC, HK1.4, Thermofisher, 1/200), Ly6G (BV510, RUO, Biolegend, 1/100), Siglec F (PE CF594, RUO, BD 1/100), F4/80 (alexa 700, clone BM8, Ozyme, 1/100), in addition to a lineage cocktail in PeCy7 (B220 clone RA3-6B2 Biolegend, CD3, clone 17A2 Biolegend, CD11c clone N418, ebioscience, NK1.1 clone PK136 Biolegend, Ter119 clone TER-119 BD Biosciences) and DAPI (1/1000) as live/dead marker. All cells were analyzed on a Ze5 (bio-rad) plate-reader cytometers. Data analysis was performed using FlowJo v10.2 software (TreeStar) and Prism v9.

### DRAG induction in mammary gland tissue

ROSACre^ERT2+/−^ DRAG^+/−^ mice were induced by one single injection of tamoxifen (0.1mg/g body weight) at P21. Mammary glands were collected 1 month after induction. Single cell dissociation was performed through enzymatic digestion (5mg/ml collagenase (Roche, 57981821) and 200U/ml hyaluronidase (Sigma, H3884) for 1h30 at 37°C under agitation. Subsequently, cells were treated with trypsin for 1 min and DNAse I and dispase for 5min at 37°C. Cell suspension was filtered through a 40µm cell strainer, and cells were stained in FACS buffer (PBS, EDTA 5mM, BSA 1%, FBS 1%) using a ‘lineage cocktail’ in APC (CD45 clone 30-F11, CD31 clone MEC13.3, Ter119+ clone TER-119, all diluted 1/100), PE EpCAM (clone G8.8, 1/100), APC/Cy7 CD49f (clone GoH3, 1/100) and DAPI. All antibodies were purchased from Biolegend. Cells were analyzed on a FACSAria^TM^ flow cytometer (BD Biosciences), and results were analysed using FlowJo software.

### DRAG induction in brain

CagGCre-ER^TM+/−^ DRAG^+/−^ mice aged 15 weeks for the uninduced group and 37 weeks for the induced group were sacrificed and their head fixed in 4% formaldehyde in PBS. Tamoxifen induction was performed in 17 week old mice as described in the tamoxifen induction section. Brains were sectioned into 30-um sections using a Leica vibratome, and sections were mounted on glass slides. Three sections, from the same rostro-caudal level in each mouse, were analyzed per mouse. Sections were imaged using a Zeiss LSM 700 confocal microscope on Tile Scan mode, using a 20x objective and 3-µm optical sections. All microscope settings (e.g., laser, gain, offset, pinhole, averaging) were kept constant for each image.

### Barcode Preprocessing and Filtering

Each recombined sequence includes nucleotide additions and deletions (referred to as the ‘barcode’) and constant parts that flank both sides of this barcode. Moreover, each barcode was associated with a random unique molecular identifier (UMI) of 12bp during the tagging PCR step.

#### Preprocessing

We use the pipeline described below to demultiplex fastq files and identify the reads that match a potential recombination of the DRAG construct. First the bcl2fastq (Illumina) program is used to demultiplex the fastq files based on the i7 index sequence. Only records that match the i7 index perfectly are considered for the next step. In the constant part of the V and the J, the reads tend to be error-prone and a consensus sequence with Ns is manually created. The Xcalibr program (https://github.com/NKI-GCF/xcalibr) is then used to extract counts for all combinations of the 12bp UMI and the recombined barcodes only for the reads that contain the constant sequence of the V (ctcgaggtcatcgaagtatcaag) at the expected coordinates. After this, the J constant part (tagcaagctcgagagtagacctactggaatcagaccgccaccatggtgagc) is aligned to the barcode part using the NBCI blast2 program ^4^. When a suitable match is found, the barcode is trimmed at the start coordinate of the match, resulting in the final matrix.

#### Barcode filtering

We used the steps described below to identify barcode sequences and remove PCR and deep-sequencing errors. First, we removed any barcode and associated UMI containing one or multiple ‘N’ values (within either the barcode, constant flanking parts or UMI). Second, barcodes that did not have an exact match to the expected constant parts (for the V region : cctcgaggtcatcgaagtatcaag and the J region : tagcaagctcgagagtagacctactggaatcaga) of the V and J that precede or follow the barcode were removed. Third, when multiple sequences were found associated with a single UMI, only the most frequently occurring barcode associated with that UMI was kept. Note that in theory, the number of different UMIs (4^12^) should be in excess to the number of template molecules present in a PCR pre-amplified sample at the point at which tagging takes place (with a maximum of 5.10^4^ expected molecules). However, we did observe rare cases in which the same UMI was associated with multiple true barcodes, presumably due to a lower than expected diversity of the tagging primer and a biased composition. Dominant barcodes that were associated with a UMI are highly likely to also be the dominant barcode associated with another UMI when samples are sequenced sufficiently deep and therefore likely to be true barcodes. Fourth, we removed UMI-barcode combinations with a read count of 10 or below to remove low-abundant combinations from our main list of barcodes. As a fifth step, we summed up the read counts for all UMI associated with the set of remaining barcodes, including the UMI from UMI-barcode combinations below the 10-read threshold for which the barcode matched one of the barcodes that passed the 10-read threshold.

To verify that the barcodes obtained match the expected structure of a VDJ recombination product, we developed an algorithm to compare barcodes to the original VDJ template and identify which nucleotides were deleted due to exonuclease activity and which ones were inserted due to Tdt activity. This enabled us to recognize barcodes containing residual error, and also to quantify barcode creation patterns (i.e. the number of deleted and inserted nucleotides at the junctions between V, D and J segments per barcode). Specifically, the algorithm performed several matching steps to the following template parts: V element=TCCAGTAG, forward D element=TCTACTATCGTTACGAC, reverse D element=GTCGTAACGATAGTAGA, and J element=GTAGCTACTACCG. The locations in between the V and D, as well as the D and J elements, are the sites where recombination can occur. Note that the residues matching the V and J region were actually longer but we did not observe deletions extending beyond the above-described residues. The algorithm started by performing exact matching to the V element, comparing residues from left to right, and to the J element from right to left. This resulted in matched V and J parts and a ‘middle part’ that contains a part of the forward or reverse D element (provided that this was not completely deleted during recombination). In order to find the most likely match of the middle part to the D element, we separately searched for the longest matches to both the forward and the reverse D element, while considering that there could be residual sequencing error within this constant element. We achieved this by starting with an attempt to match to the longest possible sequence (i.e., the entire D element of length 17 nucleotides), and decreasing the attempted match length by 1 until a match could be established. In this matching process we first searched for exact matches amongst all permutations of the considered length. For example, a comparison of the remaining middle part to a part of the D element of length 15 involves comparing to three potential D parts, i.e., a D part where two nucleotides are deleted on the left side, one where one nucleotide is deleted on both sides, and one where two nucleotides are deleted on the right side. In case no exact match could be found for the considered length, we searched for potential approximate matches in which a mismatch of a single nucleotide was allowed, provided that the mismatch did not occur in one of the two flanking residues on the left and right side (note that this implies that the minimal D fragment length for which such a mismatch can be detected is a length of 5 nucleotides). This is because a mismatch close to the flanking regions may easily be caused by nucleotide deletions and insertions during the recombination process, whereas single mismatches at locations further away from the D element flanks are more likely to be due to sequencing errors.

Having established the longest match to both the forward and the reverse D element, the longest of these two was selected for further analysis, provided that the match length was at least five nucleotides. When this longest match contained a nucleotide mismatch, the sequence was no longer considered as real and thus discarded. However, when an exact match was observed, the remaining left and right flanks of the middle part were considered insertions between V and D, and between D and J, respectively. The nucleotides of the original forward or reverse D template that were no longer present in such barcodes were considered to be deleted. For a case with a longest match of at most four nucleotides, we considered it most likely that the entire D template had been erased during recombination. In that special case, the remaining few nucleotides were assigned as follows: (i) to insertions between the V and D element in case the already recovered V element was empty and the recovered J element was not empty (because we wanted to consider the possibility that the V element was in fact non-empty but contained residual sequencing error; see below), (ii) to insertions between the D and J element in case the already recovered V and J elements were both empty (in which case we considered it likely that in fact the recovered J element was non-empty because of the large number of residues that would have been deleted on that flank otherwise), and (iii) to insertions on both insertion flanks that were equally divided amongst the V/D and D/J flank for an even number of nucleotides and with one insertion more for the D/J flank for an uneven number.

The above part of the algorithm only considered potential residual sequencing errors within the D element and not within the V and J constant elements, which was done subsequently. This was achieved by considering whether extension of the earlier detected exact matches to the V and J templates into the determined insertions between V/D and D/J, respectively, would lead to longer matches when allowing for a single mismatch in either of the constant regions. In the case of a mismatch in the constant regions, the mismatch was allowed to occur at the nucleotide immediately close to the already detected V and J parts (in the rightward and leftward direction, respectively). When a second mismatch was detected within either the second or the third nucleotide flanking the already determined exact match, an extension was not accepted. In that case, the deletions on the right side of the original V template and on the left side of the J template were determined based on the missing nucleotides. However, if the two nucleotides (in second and third position away from the exact match) did match to the original template, the position immediately flanking the earlier detected exact match was considered as an error and in that case the sequence was discarded. In summary, the algorithm detected residual sequencing errors within the constant V, D and J elements, and it determined both the insertions between V/D and D/J and the deletions from the original V, D and J templates. Note that the algorithm detected only a limited number of spurious sequences, because most of those were already removed by the other steps applied to the cellular barcoding data.

### Probability Generation Model

We used the barcode sequence lists from the previous filtering step to infer the properties of the recombination process that produce these barcodes using the IGoR algorithm, similarly to previous work ^3^. To adapt IGoR to fit the DRAG system, the genomic templates for recombination were redefined as the V, J, and D genes in the DRAG construct, adding also the inverted form of the D segment. Then, IGoR was run using all unique barcode as inputs to infer the probabilities of each possible insertion (ins) and deletion (del) scenario.

The inferred probability of recombination of a barcode σ is

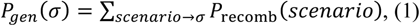

i.e. the sum of the probabilities *P*_recomb_ (*scenario*) of all recombination scenarios leading to barcode σ. Scenario probabilities are in turn given by:

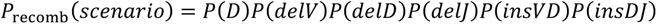

where P(D) corresponds to the probability the usage of the D or the inverted D; P(delV), P (ins VD) and P (ins DJ) correspond to insertion between the V and D segments and D and J segments respectively; P(delV), P(delD) and P(delJ) correspond to the deletion in the V,D or J segment respectively.

P(D) is calculated from the occurrence of the inverted and non-inverted form in the data, with P(D non inverted)=0.88 and P(D inverted)= 0.12. The probabilities for the insertion (P(ins)) depend both on the length of the segment (lenVD) and on its composition through a Markov Model:

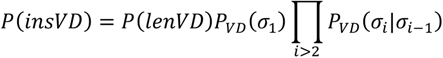

where the product runs over the non-templated inserted nucleotides. *P*(*insDJ*) is defined similarly. The inferred parameters are summarized in the Table S7–9. In the Markov model, the insertions are parameterized both by their length, and by the probability of insertion of each of the four bases, given what was the last insertion.

Note that deletion numbers for the V and J segments include the possibility of short palindromic insertions, which are given by negative deletions. Negative deletion means that, instead of being deleted, the sequence gets up to 4 additional nucleotides that are reverse-complement to the last ones. Since the J segment is longer, it can have more deletions. The D segment can be deleted from both sides, and these are correlated, so the model incorporates a joint deletions distribution, different for the reverse D. The inferred probabilities for P(del) are summarized in the Table S4–6.

This inferred model was used to calculate a generation probability (P_gen_) for every barcode using Eq. (1). The generation probability of each barcode determines how probable it is to find two cells with the same barcode, coming from different recombination events. Where indicated, we discarded barcodes above a certain threshold generation probability (P_gen_) to eliminate barcodes that are likely to be independently generated in more than one cell.

### Barcode Analysis

All analyses were carried out using R software ^5^. After running the barcode filtering pipeline, the data was placed in a count matrix for each barcode in rows and samples in columns. All barcodes that had a read value below 0.003% of total reads were set to zero to clean residual errors. The reads per sample were then renormalized to 1 or to cell numbers obtained from sorting. This renormalized matrix was used for diversity analysis. The chao index was computed using a custom script on the renormalized read to cell numbers per duplicate for each sample, using the formula below:

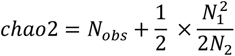

where N_obs_ is the number of barcodes observed in both duplicates, N_1_ the number of barcodes present in one duplicate and N_2_ the number of barcodes shared between duplicates. As there are different ways to compute diversity from occurrence data, we compare the results for different indexes: the bias-corrected chao2 (chao2corr), the first order jackknife (jack1), and the bootstrap (boot) (Fig. S4B) using the vegan package ^6^. The formulas for computing theses indexes are below:

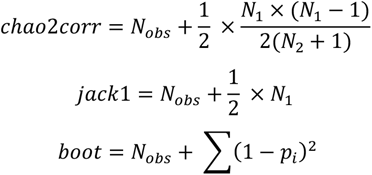

where N_obs_, N_1_ and N_2_ are defined as before and p_i_ is the frequency of the barcodes merging both duplicates.

The absolute number of HSC per blood sample was extrapolated using the chao2 index and the percentage of GFP+ cells in the sort sample (as all the bone marrow sample was sorted). We estimated that between 100-200µl of blood was collected at each time point. For the evolution of HSC diversity over time, Renyi indices were computed using a custom script on renormalized read to cell numbers to 1 per duplicate for each sample, using the formula below^7^:

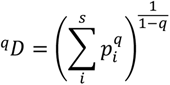

where q is the order of the diversity index, p_i_ is the frequency of a given barcode in the sample. q=0 is the richness in the sample, the number of barcodes present, q=1 is the Shannon index, q=2 is the Simpson index. For the Shannon index q=1, the limit of the ^q^D formula gives:

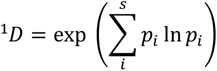

Renyi indexes were then analyzed using a gamma generalized linear mixed model, as described below.

For heatmap analyses (Fig. 3B, 4B, S4A), barcodes not present in both duplicates were removed, the technical duplicates were summed, renormalized to the arbitrary value of 10^5^ for visualization, and transformed using the hyperbolic arcsine function. Where applicable, barcodes with a P_gen_>10^x^ were filtered out. Heatmaps were generated using the heatmap package gplots ^8^, using Euclidean distance and complete linkage.

To classify barcodes into LSK, MP, and M categories (Fig. 3C), we used a previously described hand tailored classifier ^9, 10^. In summary, barcodes were classified into categories based on their presence or absence in the given cell type (LSK, MP, or M). The contribution of the sum of all barcodes in each category was computed and is displayed in figure 3C.

For the analysis of barcode sharing between duplicates, time points and mice, the Jaccard index was computed using the biomod2 ^11^ and the ade4 ^12^ packages, transformed into a fraction of barcode shared (1-jaccard^2^) and then plotted using the corrplot package ^13^.

FACS data were analyzed using FlowJo software (*Becton Dickinson)*.

### Gamma generalized linear mixed model for diversity over time

Since the Renyi entropy indices are positive and continuous variables, gamma generalized linear mixed models were fitted to the data. Let *Y_ijk_* be a random variable representing the richness measured on mouse *i*, month *j*, and subsample *k*. The conditional distribution *Y_ijk_*|*m_i_*, *S_ij_* ~ Gamma(*μ_ijk_*, *ϕ*) was assumed, with 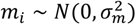 random mouse effects and 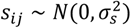 random sample effects, and *ϕ* the dispersion parameter. These random effects were included to model the correlation between the observations taken on the same mouse and duplicates. The mean was modelled with an identity link, and a piecewise-linear predictor over time (months) was used, i.e.

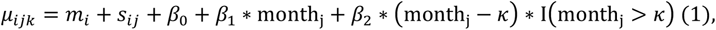

where κ is the break point estimated by maximising the profile log-likelihood of the model, and I(month_j_ > κ) is a dummy variable assuming value 1 when month_j_ > κ and 0 otherwise. Maximum likelihood estimates were obtained using the Laplace approximation for the integrals in the log-likelihood function. The best fit parameter estimates are summarized in Table S10. Goodness-of-fit of the models were assessed using half-normal plots with simulation envelopes ^14^. The models were fitted using package lme4 ^15^ from the R software.

### Gamma generalized linear mixed model for cell output per barcode over time

The cell output data consisted of continuous, strictly positive data, and therefore gamma generalized linear mixed models were used for this analysis, including random intercepts and slopes over time per mouse, and different dispersion per mouse. Let *Y_ijk_* be the response for the *i*-th mouse, *j*-th tag and *k*-th time point. It was assumed that *Y_ijk_*|*m*_0*i*_, *m*_1*i*_, *t*_0*ij*_ ~ Gamma(*μ_ijk_*, *ϕ_i_*), with 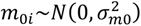 and 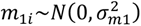 the random intercepts and slopes per mouse, respectively, 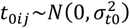 the random intercept for tag *j* within mouse *i*, and *ϕ_i_* the mouse-specific dispersion parameter. These random effects were included to model the correlation between the observations taken on the same mouse and tag. The mean was modelled with a log-link, such that

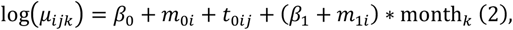

and the dispersion was also modelled using a log-link, and included different intercepts per mouse, i.e.

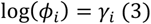

Maximum likelihood estimates were obtained using the Laplace approximation for the integrals in the log-likelihood function. Two statistical hypotheses were tested: (1) *H*_0_: β_1_ = 0 versus the alternative *H_a_*: β_1_ ≠ 0, which is equivalent to testing whether there was a trend over time, and (2) 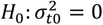 versus the alternative 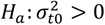, which is equivalent to testing whether within a mouse at a given time point all barcodes are equivalent. Hypotheses were tested using likelihood-ratio tests for nested models. Goodness-of-fit of the models were assessed using half-normal plots with simulation envelopes ^14^. The models were fitted using package glmmTMB ^16^ for R software.

### Bone marrow single cell transcriptomics for the HSPC composition of Figure 2

#### Bone marrow cell preparation

At sacrifice, BM was harvested from femurs, tibias and ilia and enriched using anti-CD117 magnetic beads (Miltenyi). The c-kit^+^ fraction was stained with antibodies against CD117 (c-kit APC, clone 2B8, Biolegend) and Sca-1 (Pacific Blue, clone D7, eBioscience). Cell sorting was performed on a FACSAria^TM^ (BD Biosciences) using a 70 µm nozzle at precision 0/16/0 and high efficiency. LSK (c-Kit^+^sca1^+^) cells were sorted into GFP^+^ and GFP^−^ cells from the c-Kit-enriched bone marrow fraction. <u>10X Genomics V2 3’</u> <u>Library preparation:</u> The two sorted fractions (3,000 GFP^+^ and GFP^−^ cells) were then processed using the V.2 10X genomics protocol. cDNA amplification was performed with 11-13 PCR cycles depending on the targeted cell recovery, as per the manufacturer’s recommendations. Sequencing was performed on a NovaSeq (illumina) on paired-end (PE28-8-91). <u>Single-Cell</u> <u>RNA-seq analysis:</u> Sequencing reads were processed using the default cell-ranger pre-processing pipeline and were aligned to the mouse mm10 reference genome. Gene-expression count matrices for the 642 GFP^+^ cells and 2231 GFP^−^ cells were loaded into R and analysed using Seurat v4.0. We performed QC by visual inspection of library sizes, numbers of genes expressed and mitochondrial content per cell. Cells with less than 500 genes or with a high percentage (> 10% of mitochondrial genes) were removed from downstream analyses. Cells with numbers of genes recovered were considered as doublets and filtered from the data. In this filter 1500 genes was used as an upper limit and this threshold was defined based on outlier points from plotting UMI counts and numbers of genes detected. After filtering, our count matrix contained 2,775 cells and 13,183 genes. Data was then normalised using scTransform ^17^ for which the normalized values are Pearson residuals from regularized negative binomial regression and cellular sequencing depth is used as a covariate. In our data, we observe a batch effect between GFP^+^ and GFP^−^ LSKs, likely arising because of the parallel processing of the two samples on the 10X machine as genes showed a linear increase in expression in the GFP^+^ fraction compare to the GFP^−^ fraction but not the inverse (Fig. S3A). To correct for this batch effect, we modelled the batch effect using a negative binomial model, with model residuals representing the batch-corrected expression values. We then performed dimensionality reduction on the 2,500 variably expressed genes using first principal component analysis followed by the non-linear dimensionality reduction technique UMAP ^18^ (Fig. S3B). We then performed unsupervised Louvain clustering of the data across a range of resolution parameters and chose the resolution value that led to the most stable clustering profiles ^19^ (Fig. S3C). This approach yielded 9 distinct clusters, which were manually annotated (LT-HSCs, MPP2-5) using existing markers and transcriptomic signatures ^20, 21^ (Fig. 2B and S3D). Consistent with reports that HSPCs do not form discrete cell subsets ^22–25^ we observed that many clusters co-expressed signatures of the MPP1-5 subtypes (Figure 5C, S7A-B). In cases in which clusters could not be assigned to a single HSPC subset, we named the cluster according to the combination of the different HSC and MPP signatures that they expressed ^26^. To test for possible enrichment of GFP+ cells within a given cluster, Fisher’s exact test was used. Signature expression scores were calculated using the AddModuleScore() method of Seurat V4.

### Bone marrow single cell transcriptomics for Figure 5

#### Bone marrow cell preparation

At sacrifice, BM was harvested from femurs, tibias and ilia and enriched using anti-CD117 magnetic beads (Miltenyi). The c-kit^+^ fraction was stained with antibodies against CD117 (c-kit APC, clone 2B8, Biolegend) and Sca-1 (Pacific Blue, clone D7, eBioscience). Flow cytometry was performed on a FACSAria^TM^ (BD Biosciences) or sh800 (Sony). DRAG barcoded LSK (c-Kit^+^ Sca1^+^ GFP^+^) cells were sorted using a 70 µm nozzle at precision 0/16/0 and high efficiency. <u>10X Genomics V3 3’ Library preparation:</u> Samples were processed using the 10X genomics Chromium Single Cell 3′ v3 kit. Specifically, 1,000-16,000 cells were loaded for each experiment for a targeted recovery of 500-10,000 cells. cDNA amplification was performed with 11-13 PCR cycles depending on the targeted cell recovery, as per the manufacturer’s recommendations. Sequencing was performed on a NovaSeq (illumina) on paired-end (PE28-8-91). <u>Single-Cell RNA-seq analysis:</u> Raw sequencing reads were processed using Cellranger and reads were mapped to the mouse mm10 reference genome. During filtering, Gm, Rik, and Rp genes were filtered from the dataset. Cells with less than 500 genes per cell or with a high percentage (> 15% of mitochondrial genes) were removed from downstream analyses. Cells with a UMI count greater than 50,000 were considered as doublets and removed from the data. Following these filtering procedures, the average UMI count per cell was 11,829. The median number of genes detected per cell was 2,845, 2.9% mapped to mitochondrial genes. Cell cycle annotation using the cyclone method from the scran R package showed that 13,866 cells were in G1 phase, 2,230 cells were in G2M phase, and 682 cells were in S phase. Data normalization and integration were performed using the default Seurat v4 approach FindIntegrationAnchors() followed by IntegrateData(), and differentially expressed genes were determined using a logistic regression in Seurat on the non-integrated data using the FindMarkers() function. Pathway based analyses were performed using the enrichR package ^27, 28^. To create the aged HSPC signature we performed differential gene expression analysis between HSPCs from 6.5 month old and 19 month old mice. Genes upregulated in 19 month old mice were then aggregated into a signature using the AddModuleScore() method of Seurat. Annotation of the data was obtained by unsupervised clustering of the data followed by supervised annotation in which we mapped published signatures using the AddModuleScore() method of Seurat. The MolO LT-HSC signature was taken from ^29^, and the MPP2/3/4/5 signatures were taken from ^21^ and from ^26^. Similarly to the analysis for Figure 2, in cases in which clusters could not be assigned to a single HSPC subset, we named the cluster according to the combination of different HSC and MPP signatures that they expressed. Pseudotime analysis was performed using the destiny R package. In this diffusion map approach, the algorithm creates a pairwise cell transition probability matrix. This probability is calculated by modelling cell state transitions as a random walk, in which cells can move within a local neighbourhood specified by the parameter *σ.* The greater the overlap between the gene expression neighbourhoods of two cells, the higher the transition probability. Label transfer was performed in Seurat using the FindTransferAnchors() and TransferData() methods. Briefly, this approach involves projecting the PCA structure of the reference dataset onto the query dataset. Within this shared PCA projection paired mutual nearest neighbours (anchors) are defined for each dataset. To perform label transfer, a weight matrix is defined that defines the association between query cells and anchor cells. This matrix is then multiplied by a binary classification matrix to compute a prediction score that a query cell belongs to a certain class of reference cells. In this binary classification matrix rows correspond to the different cell classes and columns correspond to the anchors. If the reference cell in the anchor pair belongs to a certain class the matrix entry is filled with a 1, otherwise 0.

### Flow cytometry analysis and statistical testing

Data analysis was performed using FlowJo^TM^ v.10 (TreeStar). Data was then exported from FlowJo and imported in GraphPad Prism. Where indicated, a Mann-Whitney test was performed.

## Code and data availability

All data and code are available at https://github.com/TeamPerie/UrbanusCosgrove-et-al-DRAG-mouse.git

## Supplementary Tables

**Table S1:**
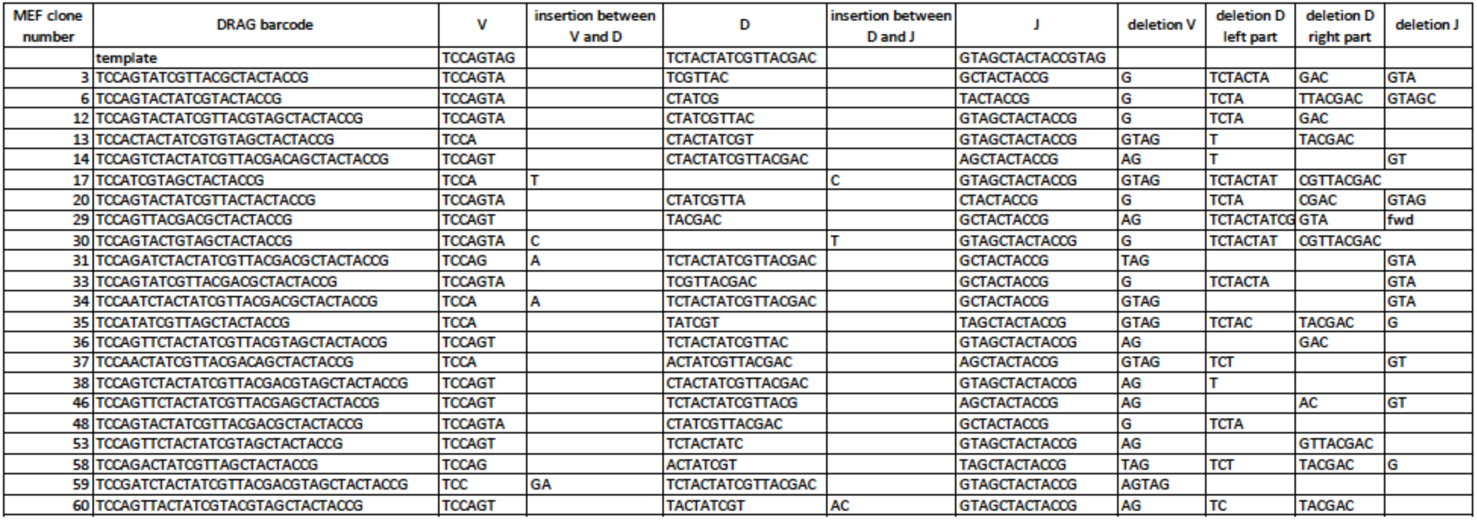
DRAG barcode sequences observed in MEF clone panel. The DRAG barcodes were split to see which parts of the sequence match the original V, D and J segments (template sequence on the first line) and compute the number of insertions and deletions.

**Table S2:**
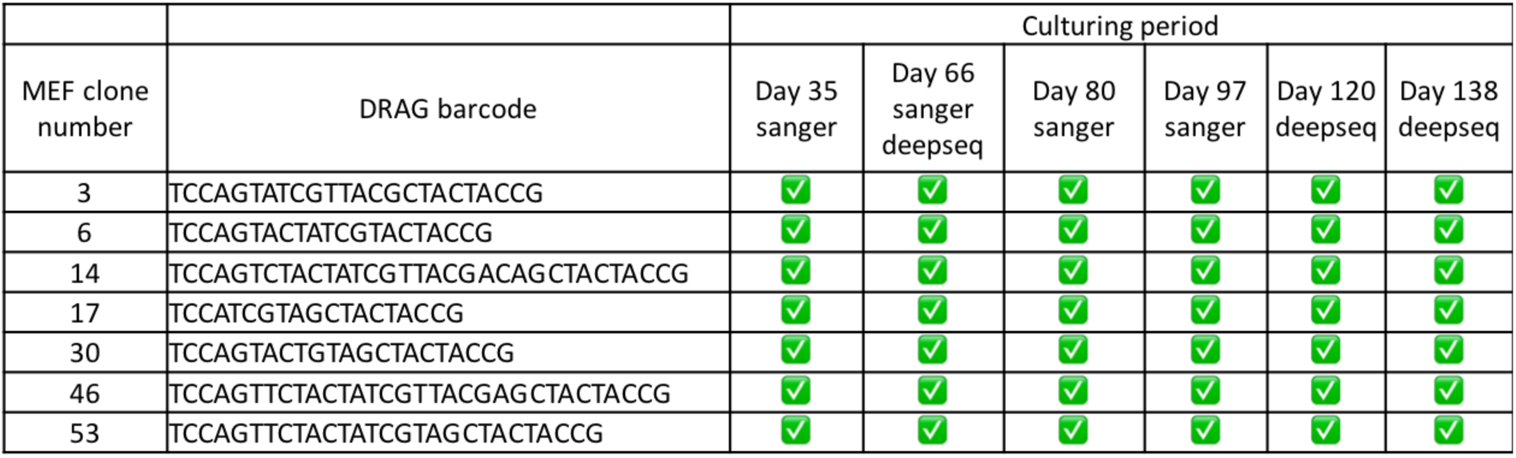
Stability of DRAG barcodes in MEF clones cultured up to 138 days. 7 MEF clones were cultured after being verified as being monoclonal and sequenced by either Sanger or Illumina NGS at the indicated time points. All clones showed the same sequence over time (indicated using a green tick).

**Table S3:** Preamp PCR cycles

| cells/half-sample | #preamp cycles |
| --- | --- |
| 300,000-600,000 | 6 |
| 150,000-300,000 | 7 |
| 75,000-150,000 | 8 |
| 35,000-75,000 | 9 |
| 17,500-35,000 | 10 |
| 9,250-17,500 | 11 |
| 4,600-9,250 | 12 |
| 2,300-4,600 | 13 |
| 1,150-2,300 | 14 |
| 575-1,150 | 15 |
| 300-575 | 16 |
| 150-300 | 17 |
| 75-150 | 18 |
| 36-75 | 19 |
| 18-36 | 20 |
| 9-18 | 21 |

**Table S4:**
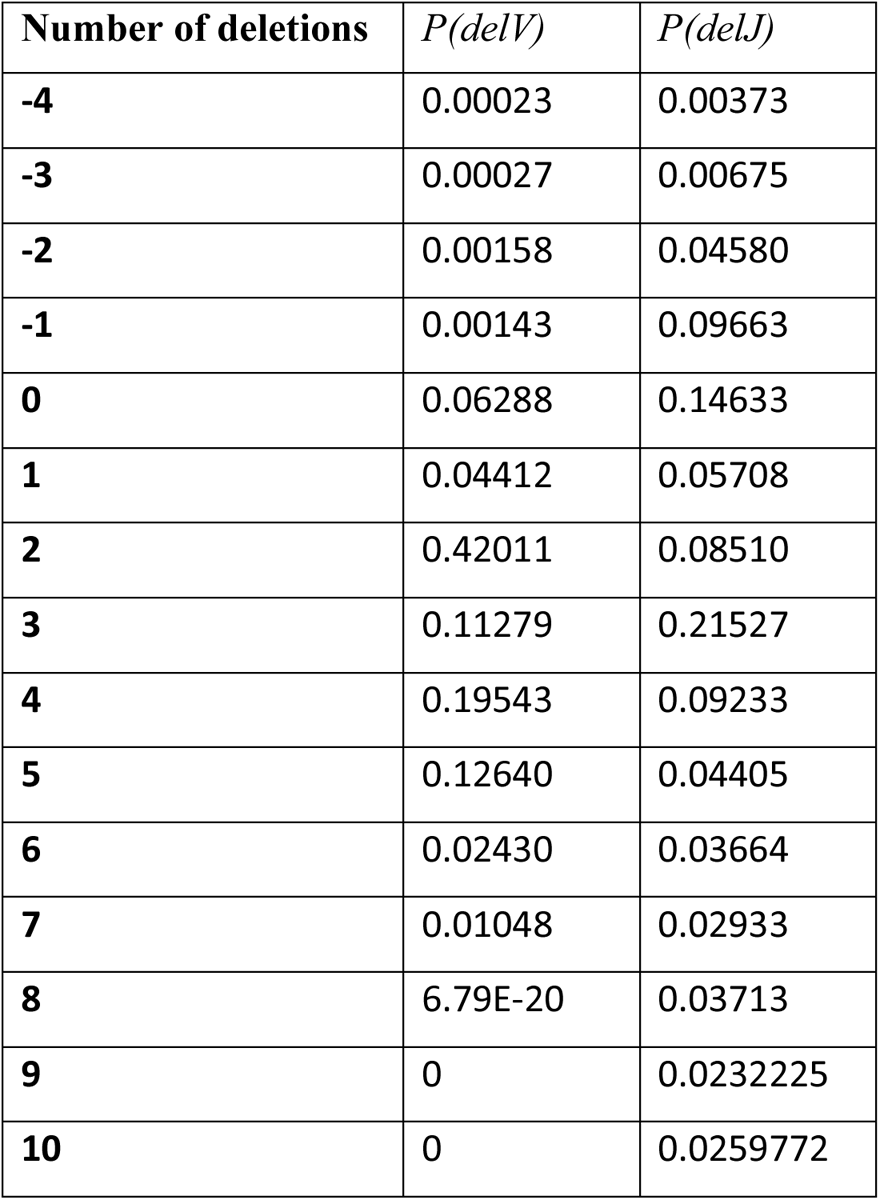
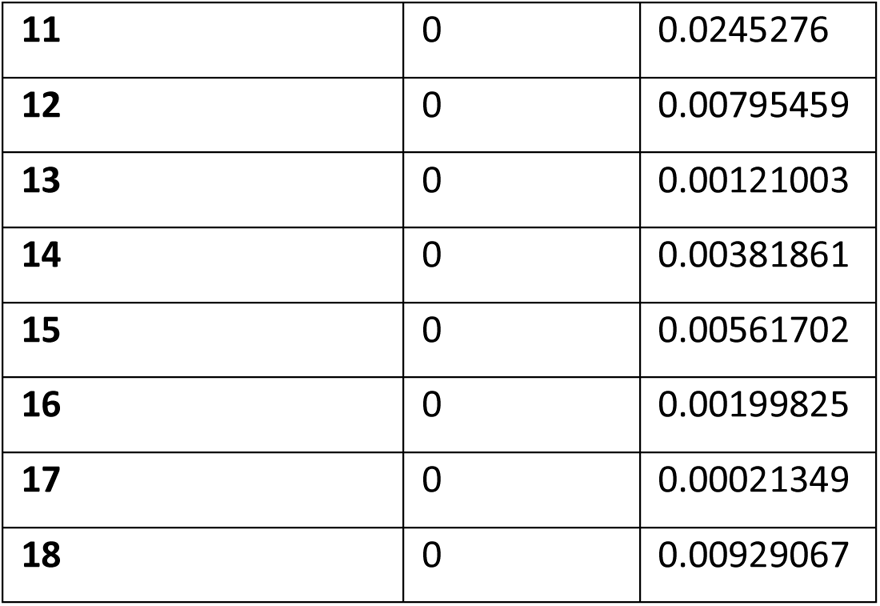
Inferred probabilities of V and J segment deletions.

| **Number of deletions** | $P(\text{del}V)$ | $P(\text{del}J)$ |
| --- | --- | --- |
| **-4** | 0.00023 | 0.00373 |
| **-3** | 0.00027 | 0.00675 |
| **-2** | 0.00158 | 0.04580 |
| **-1** | 0.00143 | 0.09663 |
| **0** | 0.06288 | 0.14633 |
| **1** | 0.04412 | 0.05708 |
| **2** | 0.42011 | 0.08510 |
| **3** | 0.11279 | 0.21527 |
| **4** | 0.19543 | 0.09233 |
| **5** | 0.12640 | 0.04405 |
| **6** | 0.02430 | 0.03664 |
| **7** | 0.01048 | 0.02933 |
| **8** | 6.79E-20 | 0.03713 |
| **9** | 0 | 0.0232225 |
| **10** | 0 | 0.0259772 |
| **11** | 0 | 0.0245276 |
| **12** | 0 | 0.00795459 |
| **13** | 0 | 0.00121003 |
| **14** | 0 | 0.00381861 |
| **15** | 0 | 0.00561702 |
| **16** | 0 | 0.00199825 |
| **17** | 0 | 0.00021349 |
| **18** | 0 | 0.00929067 |

**Table S5:** Inferred joint probabilities for D segment deletions. 5’ deletions are depicted in rows, 3’ deletions in columns

| $P(\text{delD})$ | 0 | 1 | 2 | 3 | 4 | 5 | 6 | 7 | 8 | 9 | 10 | 11 | 12 | 13 | 14 | 15 | 16 |
| --- | --- | --- | --- | --- | --- | --- | --- | --- | --- | --- | --- | --- | --- | --- | --- | --- | --- |
| 0 | 0.012 | 0.005 | 0.009 | 0.005 | 0.011 | 0.003 | 0.009 | 0.009 | 0.003 | 0.001 | 0.000 | 0.000 | 0.001 | 0.000 | 0.000 | 0.000 | 0.000 |
| 1 | 0.031 | 0.011 | 0.020 | 0.020 | 0.030 | 0.009 | 0.012 | 0.020 | 0.005 | 0.001 | 0.002 | 0.003 | 0.004 | 0.001 | 0.002 | 0.005 | 0.016 |
| 2 | 0.009 | 0.002 | 0.003 | 0.003 | 0.004 | 0.002 | 0.002 | 0.002 | 0.000 | 0.000 | 0.000 | 0.002 | 0.001 | 0.000 | 0.004 | 0.008 | 0.000 |
| 3 | 0.032 | 0.015 | 0.019 | 0.019 | 0.031 | 0.009 | 0.017 | 0.024 | 0.007 | 0.002 | 0.002 | 0.002 | 0.002 | 0.000 | 0.002 | 0.000 | 0.000 |
| 4 | 0.021 | 0.009 | 0.011 | 0.009 | 0.011 | 0.005 | 0.008 | 0.007 | 0.003 | 0.001 | 0.001 | 0.002 | 0.001 | 0.003 | 0.000 | 0.000 | 0.000 |
| 5 | 0.016 | 0.003 | 0.007 | 0.005 | 0.006 | 0.001 | 0.002 | 0.002 | 0.001 | 0.000 | 0.000 | 0.000 | 0.000 | 0.000 | 0.000 | 0.000 | 0.000 |
| 6 | 0.019 | 0.004 | 0.007 | 0.006 | 0.007 | 0.004 | 0.005 | 0.003 | 0.000 | 0.001 | 0.000 | 0.001 | 0.000 | 0.000 | 0.000 | 0.000 | 0.000 |
| 7 | 0.003 | 0.001 | 0.002 | 0.001 | 0.001 | 0.000 | 0.000 | 0.000 | 0.004 | 0.000 | 0.000 | 0.000 | 0.000 | 0.000 | 0.000 | 0.000 | 0.000 |
| 8 | 0.020 | 0.004 | 0.008 | 0.009 | 0.012 | 0.003 | 0.001 | 0.002 | 0.001 | 0.000 | 0.000 | 0.000 | 0.000 | 0.000 | 0.000 | 0.000 | 0.000 |
| 9 | 0.003 | 0.001 | 0.001 | 0.000 | 0.000 | 0.000 | 0.000 | 0.000 | 0.000 | 0.000 | 0.000 | 0.000 | 0.000 | 0.000 | 0.000 | 0.000 | 0.000 |
| 10 | 0.001 | 0.000 | 0.001 | 0.000 | 0.000 | 0.001 | 0.000 | 0.000 | 0.000 | 0.000 | 0.000 | 0.000 | 0.000 | 0.000 | 0.000 | 0.000 | 0.000 |
| 11 | 0.002 | 0.002 | 0.002 | 0.000 | 0.000 | 0.000 | 0.000 | 0.000 | 0.000 | 0.000 | 0.000 | 0.000 | 0.000 | 0.000 | 0.000 | 0.000 | 0.000 |
| 12 | 0.002 | 0.001 | 0.000 | 0.002 | 0.000 | 0.000 | 0.000 | 0.000 | 0.000 | 0.000 | 0.000 | 0.000 | 0.000 | 0.000 | 0.000 | 0.000 | 0.000 |
| 13 | 0.001 | 0.001 | 0.001 | 0.000 | 0.000 | 0.000 | 0.000 | 0.000 | 0.000 | 0.000 | 0.000 | 0.000 | 0.000 | 0.000 | 0.000 | 0.000 | 0.000 |
| 14 | 0.001 | 0.000 | 0.000 | 0.000 | 0.000 | 0.000 | 0.000 | 0.000 | 0.000 | 0.000 | 0.000 | 0.000 | 0.000 | 0.000 | 0.000 | 0.000 | 0.000 |
| 15 | 0.001 | 0.000 | 0.000 | 0.000 | 0.000 | 0.000 | 0.000 | 0.000 | 0.000 | 0.000 | 0.000 | 0.000 | 0.000 | 0.000 | 0.000 | 0.000 | 0.000 |
| 16 | 0.000 | 0.000 | 0.000 | 0.000 | 0.000 | 0.000 | 0.000 | 0.000 | 0.000 | 0.000 | 0.000 | 0.000 | 0.000 | 0.000 | 0.000 | 0.000 | 0.000 |

**Table S6:** Inferred probabilities for inverted D segment deletions. 5’ deletions are depicted in rows, 3’ deletion in columns

| $P(\text{delD})$ | 0 | 1 | 2 | 3 | 4 | 5 | 6 | 7 | 8 | 9 | 10 | 11 | 12 | 13 | 14 | 15 | 16 |
| --- | --- | --- | --- | --- | --- | --- | --- | --- | --- | --- | --- | --- | --- | --- | --- | --- | --- |
| 0 | 0.012 | 0.006 | 0.004 | 0.007 | 0.000 | 0.046 | 0.009 | 0.003 | 0.015 | 0.004 | 0.001 | 0.000 | 0.000 | 0.000 | 0.000 | 0.000 | 0.000 |
| 1 | 0.000 | 0.000 | 0.000 | 0.000 | 0.000 | 0.002 | 0.000 | 0.000 | 0.000 | 0.000 | 0.000 | 0.009 | 0.000 | 0.001 | 0.025 | 0.015 | 0.003 |
| 2 | 0.020 | 0.009 | 0.015 | 0.013 | 0.012 | 0.128 | 0.026 | 0.017 | 0.034 | 0.010 | 0.002 | 0.007 | 0.001 | 0.011 | 0.009 | 0.008 | 0.000 |
| 3 | 0.005 | 0.021 | 0.003 | 0.001 | 0.001 | 0.006 | 0.001 | 0.001 | 0.005 | 0.000 | 0.000 | 0.000 | 0.000 | 0.000 | 0.030 | 0.000 | 0.000 |
| 4 | 0.003 | 0.002 | 0.001 | 0.001 | 0.001 | 0.007 | 0.000 | 0.000 | 0.000 | 0.000 | 0.002 | 0.002 | 0.000 | 0.000 | 0.000 | 0.000 | 0.000 |
| 5 | 0.000 | 0.014 | 0.011 | 0.009 | 0.005 | 0.021 | 0.003 | 0.000 | 0.004 | 0.002 | 0.039 | 0.000 | 0.000 | 0.000 | 0.000 | 0.000 | 0.000 |
| 6 | 0.001 | 0.004 | 0.003 | 0.004 | 0.002 | 0.003 | 0.000 | 0.004 | 0.002 | 0.014 | 0.000 | 0.000 | 0.000 | 0.000 | 0.000 | 0.000 | 0.000 |
| 7 | 0.006 | 0.002 | 0.004 | 0.003 | 0.003 | 0.012 | 0.001 | 0.004 | 0.012 | 0.010 | 0.000 | 0.000 | 0.000 | 0.000 | 0.000 | 0.000 | 0.000 |
| 8 | 0.001 | 0.001 | 0.000 | 0.000 | 0.000 | 0.000 | 0.000 | 0.001 | 0.001 | 0.000 | 0.000 | 0.000 | 0.000 | 0.000 | 0.000 | 0.000 | 0.000 |
| 9 | 0.000 | 0.000 | 0.000 | 0.000 | 0.001 | 0.000 | 0.008 | 0.000 | 0.000 | 0.000 | 0.000 | 0.000 | 0.000 | 0.000 | 0.000 | 0.000 | 0.000 |
| 10 | 0.000 | 0.000 | 0.000 | 0.000 | 0.000 | 0.002 | 0.000 | 0.000 | 0.000 | 0.000 | 0.000 | 0.000 | 0.000 | 0.000 | 0.000 | 0.000 | 0.000 |
| 11 | 0.000 | 0.000 | 0.001 | 0.004 | 0.006 | 0.000 | 0.000 | 0.000 | 0.000 | 0.000 | 0.000 | 0.000 | 0.000 | 0.000 | 0.000 | 0.000 | 0.000 |
| 12 | 0.000 | 0.000 | 0.000 | 0.000 | 0.000 | 0.000 | 0.000 | 0.000 | 0.000 | 0.000 | 0.000 | 0.000 | 0.000 | 0.000 | 0.000 | 0.000 | 0.000 |
| 13 | 0.000 | 0.000 | 0.002 | 0.000 | 0.000 | 0.000 | 0.000 | 0.000 | 0.000 | 0.000 | 0.000 | 0.000 | 0.000 | 0.000 | 0.000 | 0.000 | 0.000 |
| 14 | 0.002 | 0.006 | 0.000 | 0.000 | 0.000 | 0.000 | 0.000 | 0.000 | 0.000 | 0.000 | 0.000 | 0.000 | 0.000 | 0.000 | 0.000 | 0.000 | 0.000 |
| 15 | 0.001 | 0.000 | 0.000 | 0.000 | 0.000 | 0.000 | 0.000 | 0.000 | 0.000 | 0.000 | 0.000 | 0.000 | 0.000 | 0.000 | 0.000 | 0.000 | 0.000 |
| 16 | 0.000 | 0.000 | 0.000 | 0.000 | 0.000 | 0.000 | 0.000 | 0.000 | 0.000 | 0.000 | 0.000 | 0.000 | 0.000 | 0.000 | 0.000 | 0.000 | 0.000 |

**Table S7:** Inferred parameters of the Markov model for insertions between V and D segments, with the probabilities for inserting different bases n1 in rows, given the last inserted base n2 in columns, in the 5’ direction.

| $P_{VD}(n_1\vert n_2)$ | A | C | G | T |
| --- | --- | --- | --- | --- |
| A | 0.20 | 0.10 | 0.24 | 0.26 |
| C | 0.26 | 0.51 | 0.16 | 0.39 |
| G | 0.41 | 0.13 | 0.49 | 0.16 |
| T | 0.131 | 0.25 | 0.10 | 0.18 |

**Table S8:** Inferred parameters of the Markov model for insertions between the D and J segments, with the probabilities for inserting different bases n1 in rows, given the last inserted base n2 in columns, in the 3’ direction.

| $P_{DJ}(n_1\vert n_2)$ | A | C | G | T |
| --- | --- | --- | --- | --- |
| A | 0.26 | 0.18 | 0.24 | 0.09 |
| C | 0.17 | 0.46 | 0.14 | 0.41 |
| G | 0.39 | 0.16 | 0.53 | 0.29 |
| T | 0.17 | 0.20 | 0.09 | 0.20 |

**Table S9:** Inferred probabilities for insertions

| number of insertions | $P(insVD)$ | $P(insDJ)$ |
| --- | --- | --- |
| 0 | 0.77 | 0.38 |
| 1 | 0.05 | 0.09 |
| 2 | 0.05 | 0.14 |
| 3 | 0.04 | 0.16 |
| 4 | 0.04 | 0.10 |
| 5 | 0.02 | 0.05 |
| 6 | 0.02 | 0.04 |
| 7 | 0.01 | 0.02 |
| 8 | 0.01 | 0.01 |
| 9 | 1E-03 | 0.01 |
| 10 | 3E-03 | 0.01 |
| 11 | 1.03E-03 | 1.80 E-03 |
| 12 | 3.87E-04 | 2.72 E-03 |
| 13 | 2.82E-04 | 3.94 E-04 |
| 14 | 0 | 1.03 E-03 |
| 15 | 5.49E-05 | 0 |
| 16 | 0 | 0 |
| 17 | 4.76E-05 | 0 |
| 18 | 2.46E-26 | 2.24E-04 |
| 19 | 0 | 1.27E-10 |

**Table S10:**
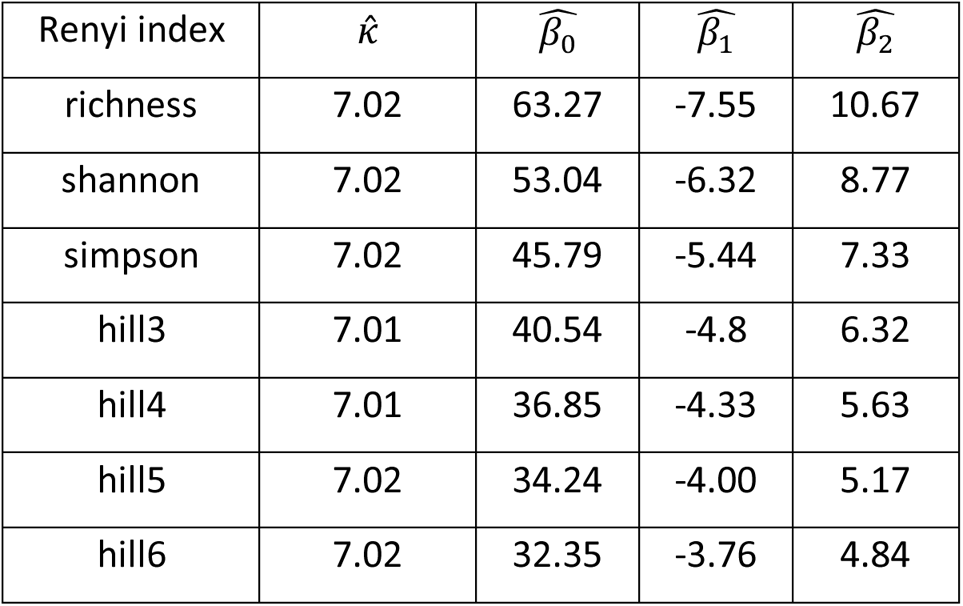
Parameter estimates from equation (1) of the gamma generalized linear mixed model for barcode diversity over time for different Renyi indexes.

Table S11: List of i7 indexes (attached)

Table S12: Comparisons between in situ barcoding methods (attached)

Table S13: Differentially expressed genes across clusters (Figure 5) (attached)

Table S14: Differentially expressed genes across HSPCs taken from mice of different ages (attached)

Table S15: Pathway enrichment analysis comparing HSPCs taken from mice of different ages (attached)

## Supplementary figures

**Figure S1:**
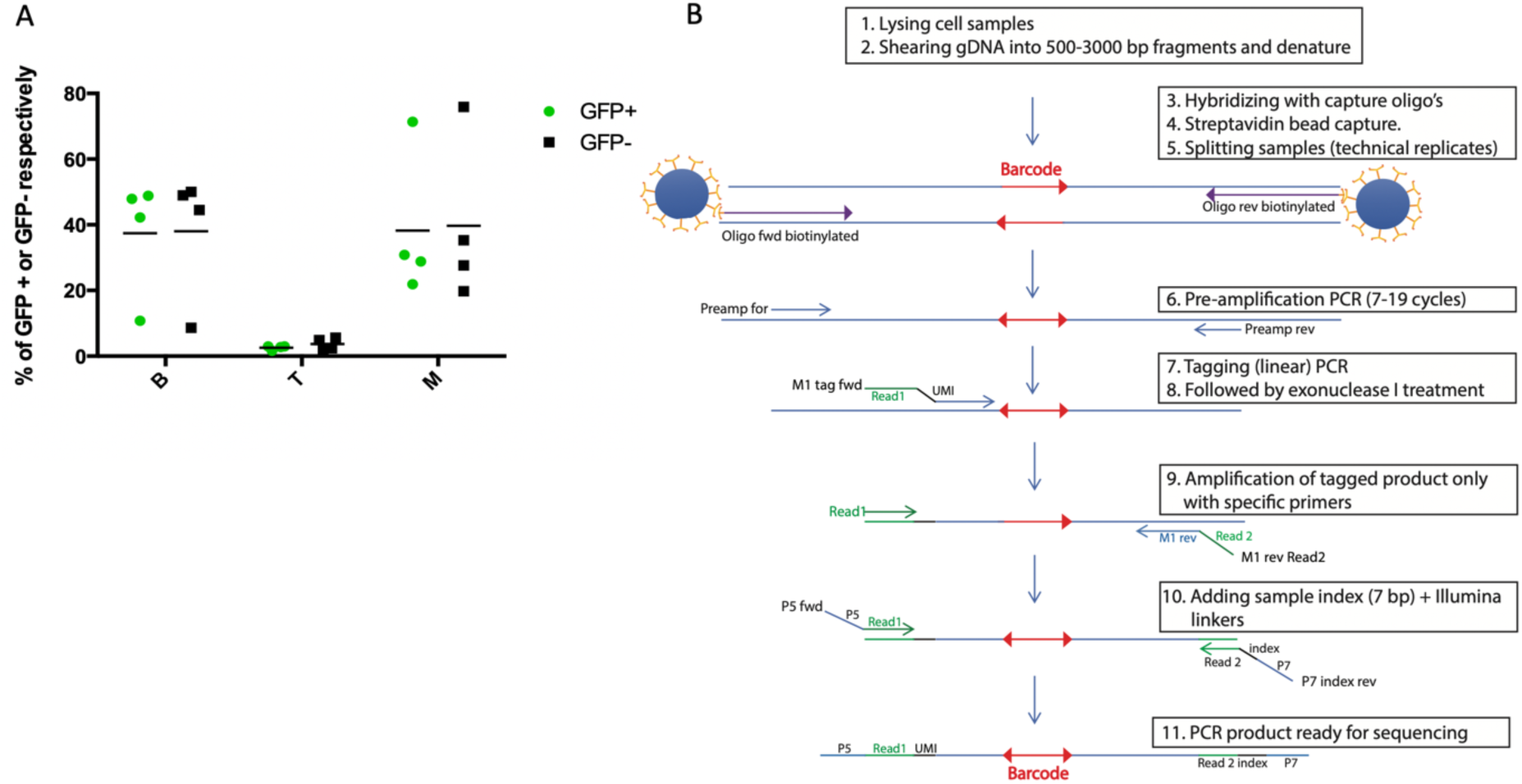
A. DRAG induction is neutral with respect to cell differentiation. Data depict the percentage of B cells (CD19^+^), T cells (CD3^+^) and myeloid cells (CD11b^+^) within either the GFP^+^ (green) or GFP^−^ (black) cell population in blood of tamoxifen-induced mice, 9 months after induction. The line represents the median and individual points the 4 mice analyzed. B. Sample processing pipeline for DRAG barcode amplification and deep sequencing.

**Figure S2:**
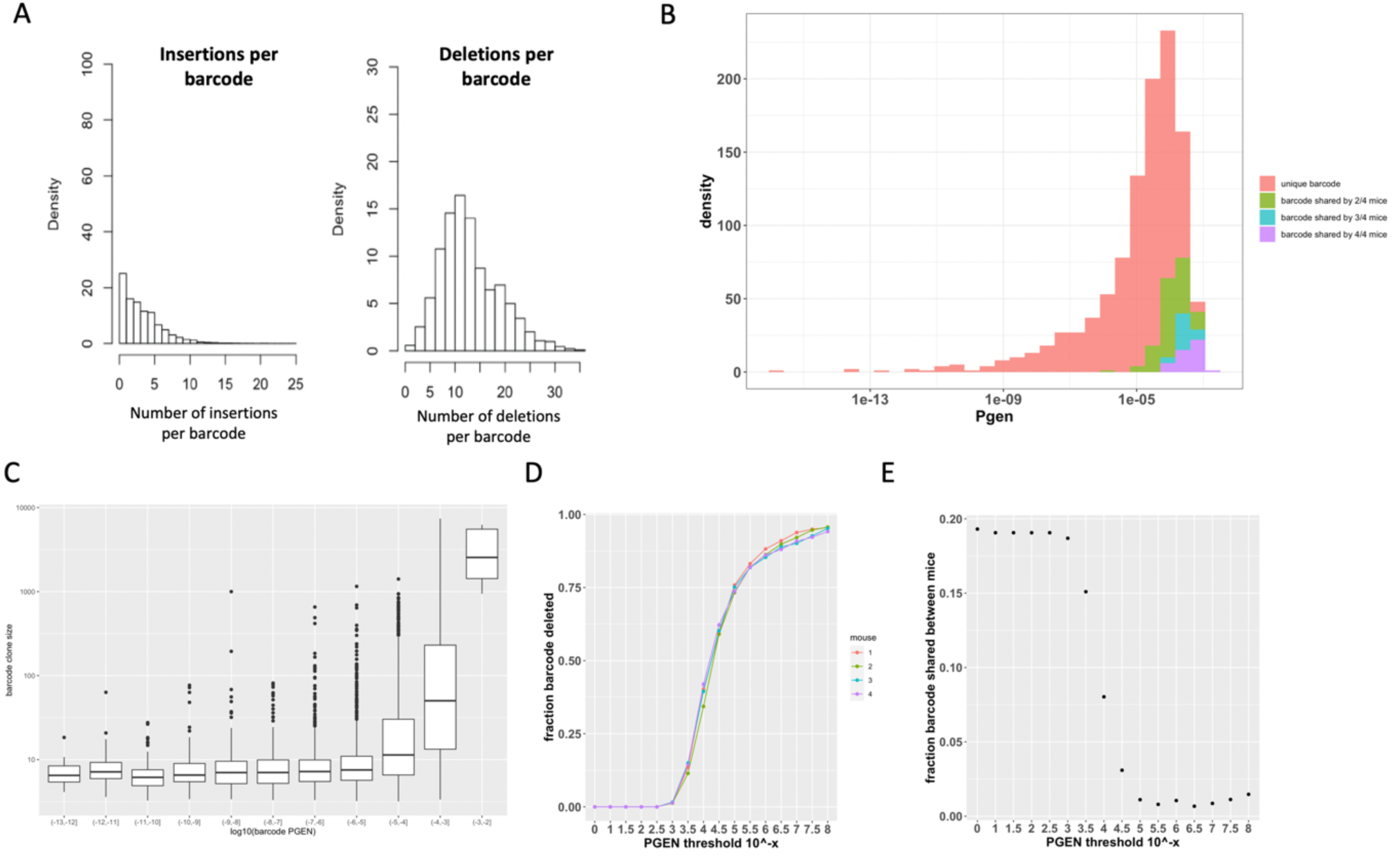
Quality control data for DRAG barcode recombination. A. Left, middle, bar graphs depicting the number of nucleotides inserted and deleted between the V, D, and J segments. All summary statistics in this graph are derived from an experiment using 4 mice. B. Distribution of the generation probability of barcodes (Pgen) of barcodes observed in an experiment using 4 mice. Pgen was calculated using the model described in the methods section. C. Frequency of reads of barcodes as a function of their estimated Pgen. D. Fraction of total barcodes discarded per mouse for different Pgen threshold values (Pgen < 10^−x^). For example, when using Pgen<10^−4^, on average 39+/−3% of barcodes are discarded. E. Fraction of barcodes that are present in more than one mouse for different values of the Pgen threshold (Pgen < 10^−x^). For example, when using Pgen<10^−4^, 92% of the retained barcodes were unique to an individual mouse.

**Figure S3:**
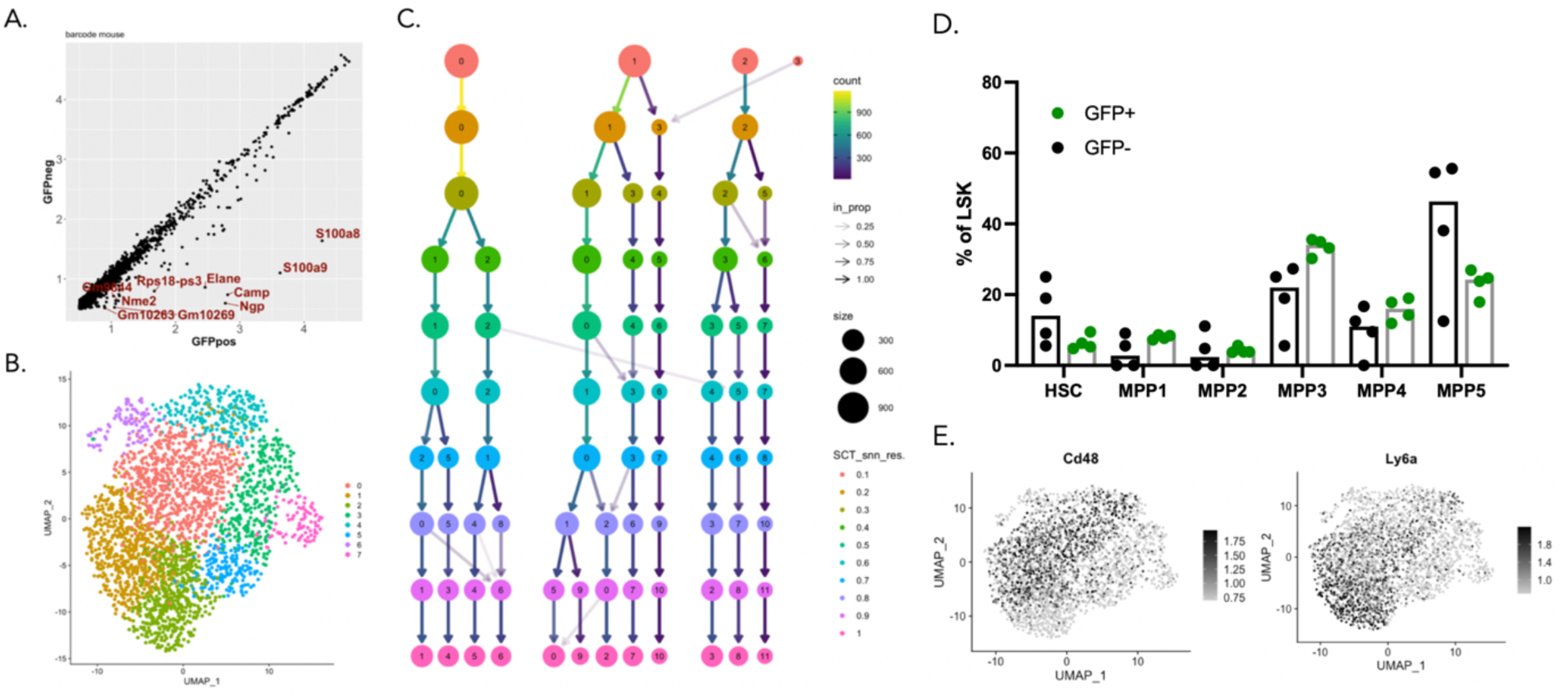
Supplementary Data for scRNAseq analysis in Figure 2. A. Comparison of the mean log expression of genes between GFP^+^ and GFP^−^ LSK cells prior to batch correction. B. Unsupervised Louvain clustering analysis of the LSK compartment using the Seurat package plotted using a UMAP representation. C. Cluster stability analysis for scRNAseq profiling of HSPCs, showing the relationship between clusters at different resolution parameters used in Seurat. The size of each node represents the number of cells in each cluster and colors represent different values of the resolution parameter in Seurats’ implementation of the Louvain clustering algorithm. D. Flow cytometry quantification of proportion of GFP+ and GFP− HSPC subsets of 6 month old mice. Each point represents 1 mouse and n = 4. No statistically significant differences between GFP− and GFP+ representation amongst the HSPC subsets were observed. Statistical significance was tested using a paired Wilcoxon-Test. Full gating strategy is provided in Figure S12A. E. Overlay of *Cd48* and *Ly6a* expression onto the UMAP representation of the data.

**Figure S4:**
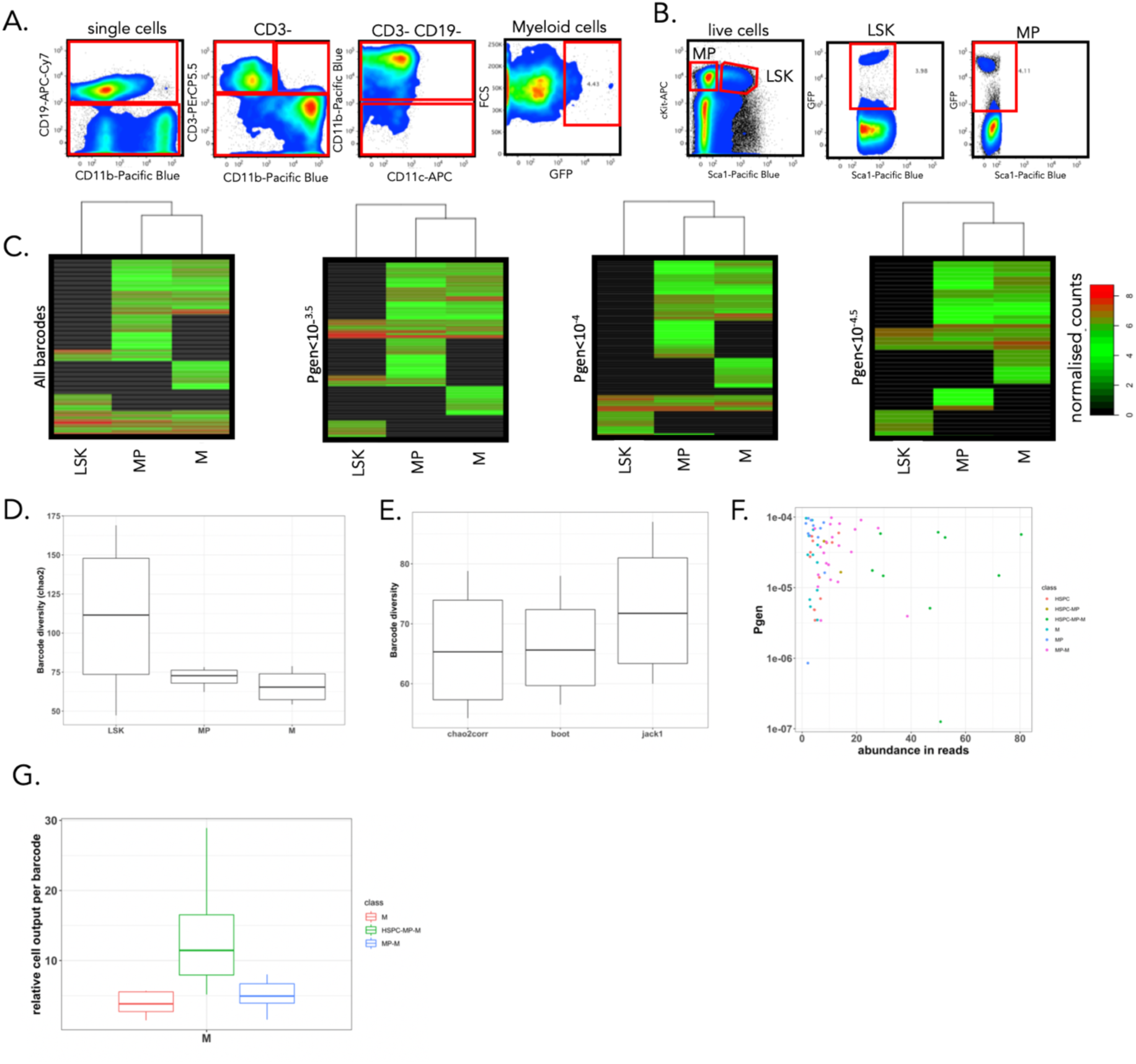
Supporting analyses for Figure 3. A. Cell sorting strategy used to obtain bone marrow myeloid cells (c-Kit enriched fraction) and blood myeloid cells. In addition, a representative plot of GFP expression in the sorted cell population is depicted. B. Cell sorting strategy for bone marrow myeloid progenitors (MP) and HSPC (LSK) from the c-Kit enriched fractions of whole bone marrow. C. Heatmap representation of barcode output in bone marrow HSPC, MP, and myeloid cells, as defined in Fig3B, at month 15 post-induction. Data are depicted for different values of the threshold for barcode generation probability Pgen (retaining 227, 164, 97, or 42 barcodes for analysis for the Pgen values depicted from left to right). Pooled data of 4 mice, renormalized, arcsine transformed data clustered by complete linkage using Euclidean distance are depicted. D. Chao2 estimate of the diversity of barcodes in bone marrow HSPC, MP, and myeloid cells in bone marrow and blood 15 months post-induction. mean and SD over 4 mice. E. Comparison of different diversity estimators based on abundance data. The estimators used are the bias-corrected chao2 (chao2corr), the first order jackknife (jack1), the bootstrap (boot). F. Probability of DRAG barcode generation as a function of read abundance, color coded by the different classes of output as in Figure 3C and 3D. G. The relative cell output represents the fraction of reads per barcode of the total reads found in myeloid cells. This relative cell output is presented per barcode category as defined in Figure 3C and 3D. The mean and SD over barcodes obtained from 4 mice is displayed. The colors represent the barcode categories as in Figure 3C and 3D.

**Figure S5:**
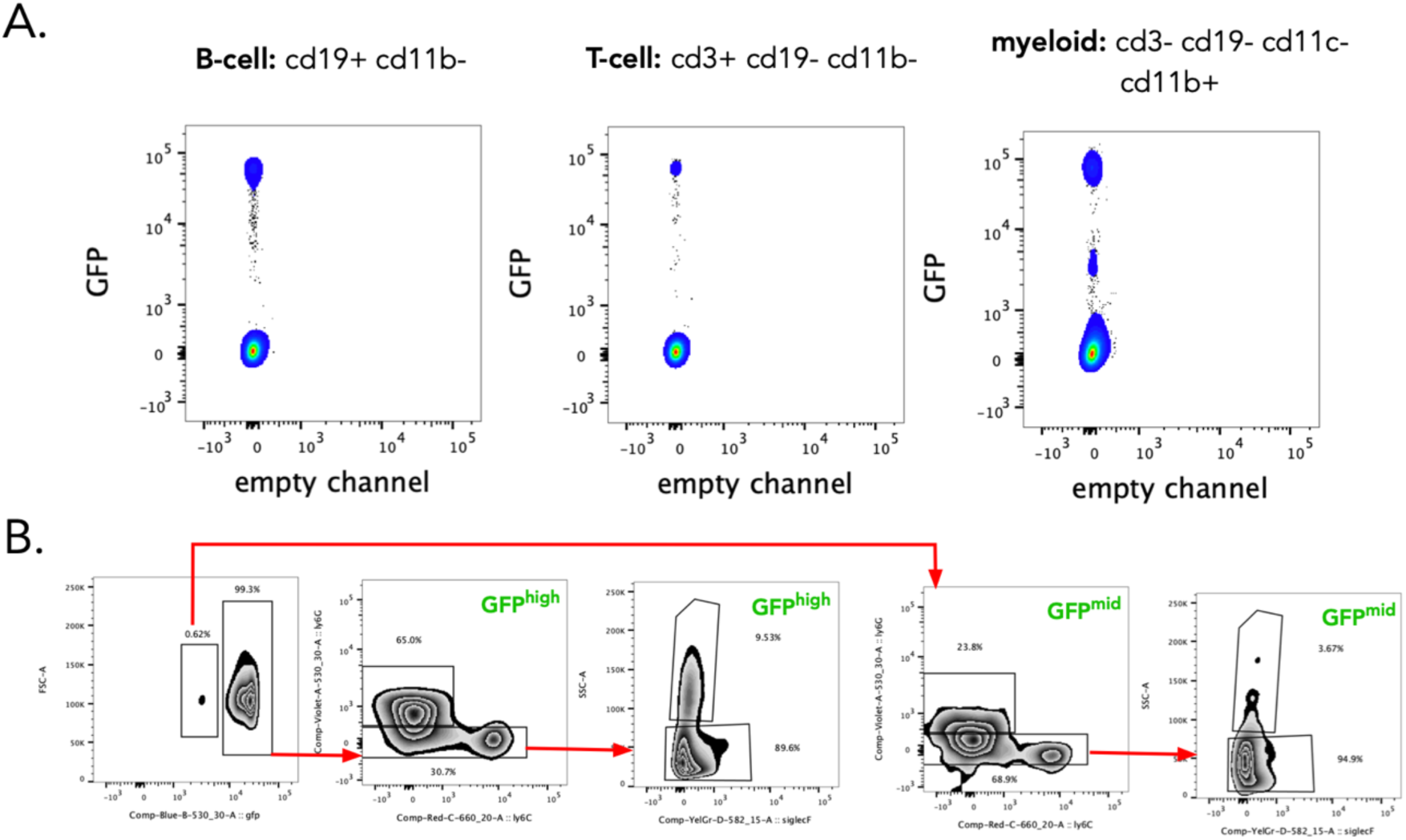
Analysis of GFP expression levels upon DRAG recombination in different cell compartments. (A) DRAG induced GFP expression within lymphoid and myeloid lineages. Only within the myeloid lineage a separate GFP^mid^ population is observed. (B) surface marker expression patterns for GFP^mid^ and GFP^high^ myeloid cell populations.

**Figure S6:**
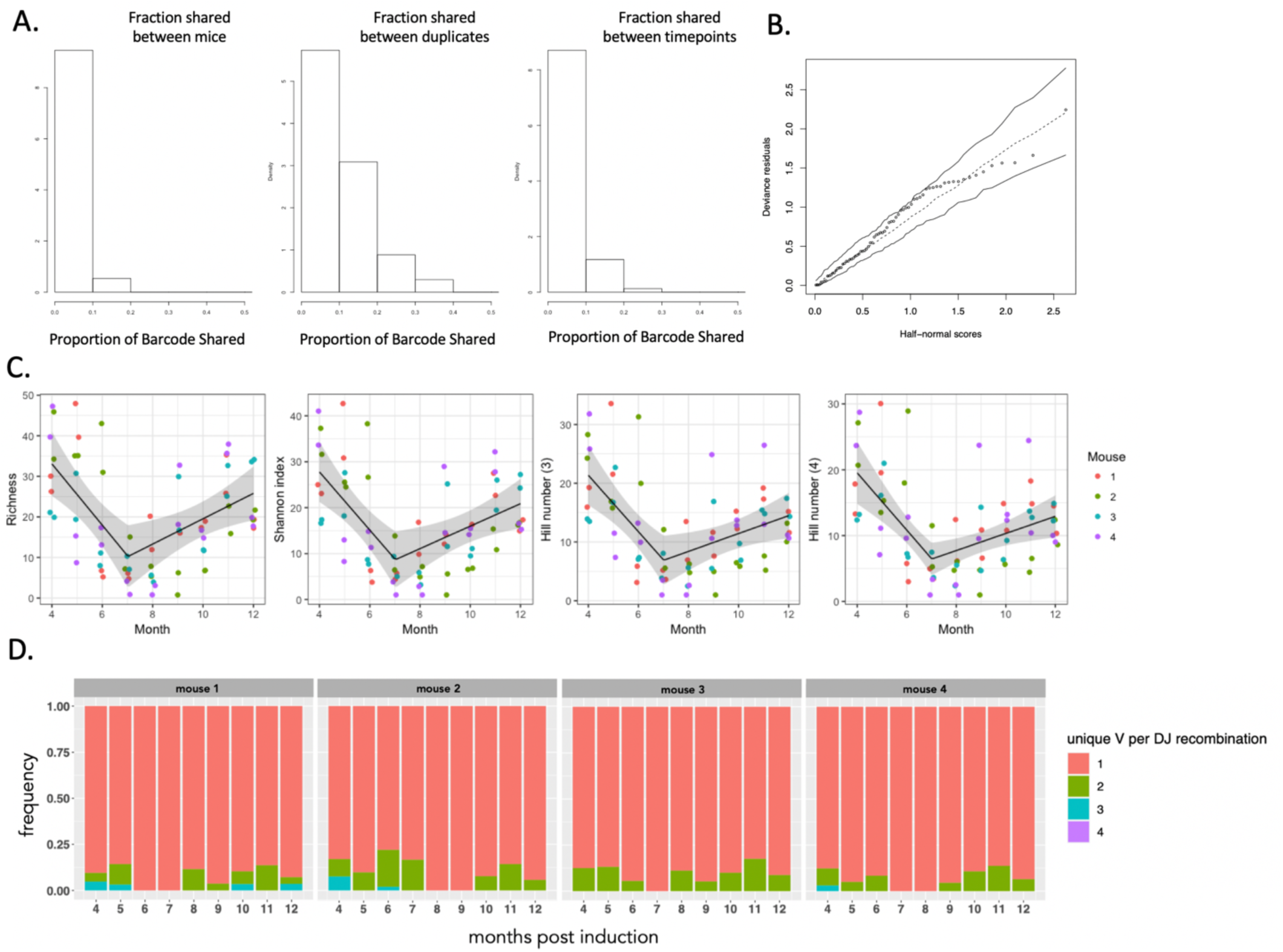
Supporting analyses for Figure 3. A. Fraction of barcodes in myeloid cells shared between mice (indicative of frequently occurring barcodes), between duplicates (indicative of sampling efficiency at a given time point) and between time points. Pooled data of four mice for the experimental set up shown in figure 3A. B. Half-normal plot of the conditional deviance residuals for the generalized linear mixed model (GLMM) used in Figure 3G including a simulated envelope. The simulated envelope (solid lines) is obtained by simulating 99 response variables assuming the fitted model is the true model, refitting the same GLMM to the simulated samples, then obtaining and sorting the conditional deviance residuals in absolute value, and calculating the 2.5% and 97.5% percentiles for each order statistic, while the dashed line represents the median. The envelope is such that for a well-fitted model, most points are expected to fall within the envelope. In this case, all 72 points lie within the simulated envelope. C. Four different diversity estimates (Richness, Shannon index, Hill 3 and 4 number) of the barcodes in the myeloid cells in blood between 4 months and 12 months per mouse. Each sample was measured in duplicate. The black line is the best fitted value of the gamma generalized linear mixed model with a break point. The grey ribbon represents the 95% CI for the true means. D. For each D and J recombination, the number of associated V regions was computed across all barcodes. The % of total barcodes (recombination) that had one, two or more V regions associated with one DJ recombination is plotted for the indicated times after barcode induction. The color represents the number of associated V regions and each of the four graph displays the result from one of four mice.

**Figure S7:**
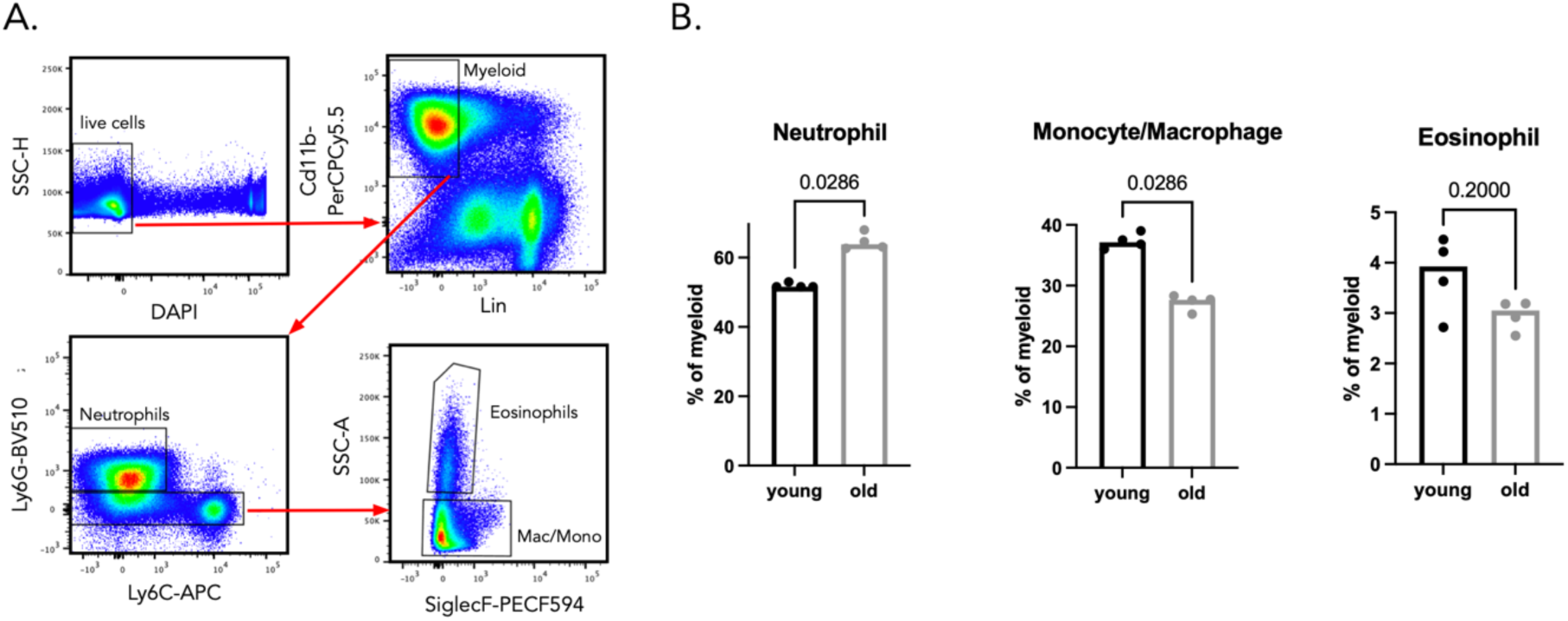
Changes in the composition of the bone marrow myeloid compartment during ageing. A. Gating strategy to identify neutrophils, eosinophils and monocytes-macrophages. B. Percentage of the indicated cell types in the bone marrow myeloid population of young and old mice. N = 4 mice per group, each point represents 1 mouse and statistical comparisons were made using a Mann-Whitney test. Young mice were 6.5 months old and old mice were 19 months old at the time of sample processing. Barcodes were induced at 10-20 weeks after birth.

**Figure S8:**
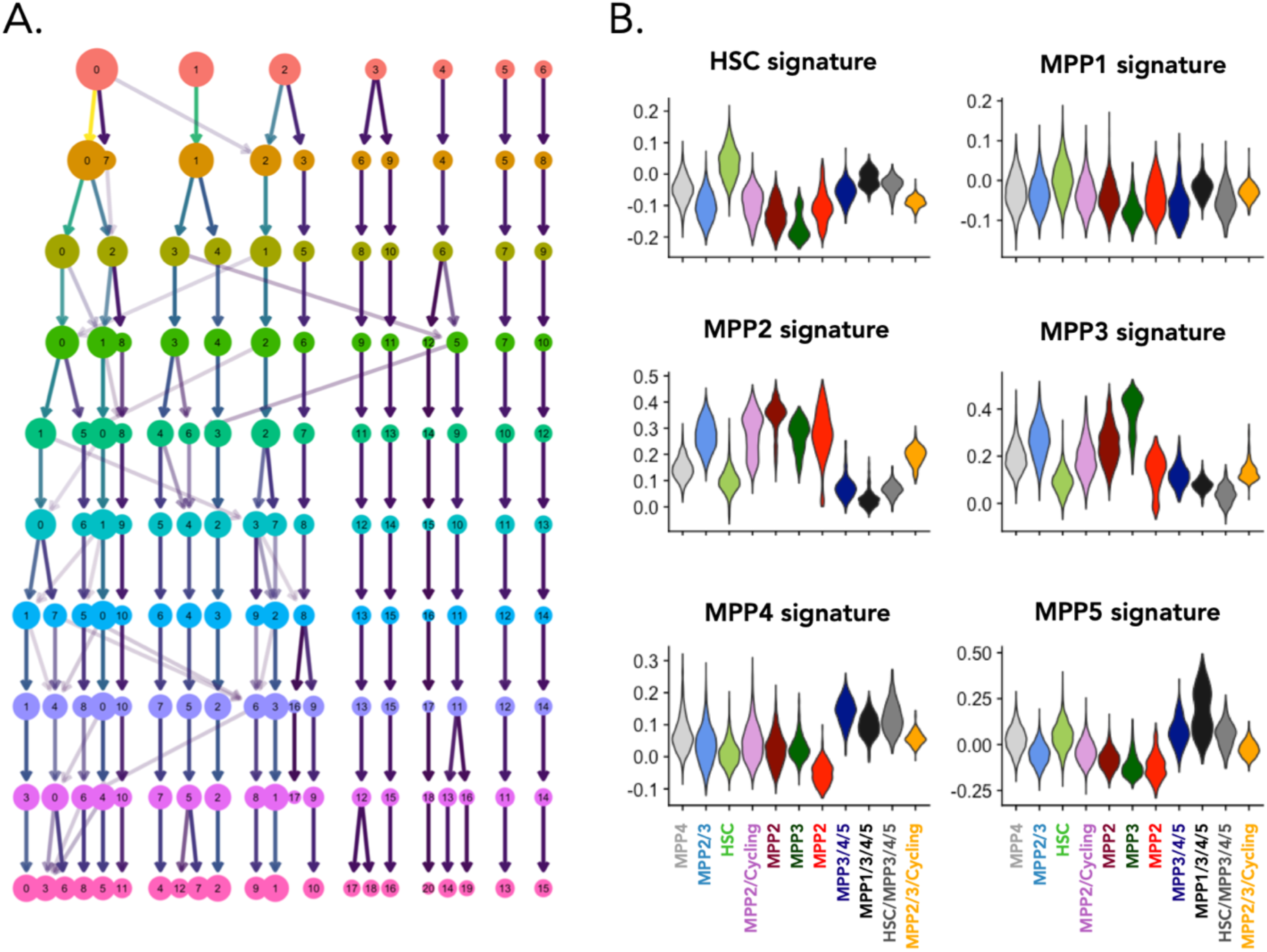
Supporting analyses for scRNAseq profiling of aged HSPCs. (A) Cluster stability analysis. Each row represents a different resolution parameter of the Seurat default clustering algorithm. Each node represents a cluster and arrows represent the relationship between clusters across different resolution parameters. The size of each node is scaled to the number of cells in the respective cluster. (B) Expression of HSC and MPP signatures from Sommerkamp et al (2021) amongst different clusters. Signature expression for each cell was calculated by taking the mean expression across all genes (after background correction)

**Figure S9:**
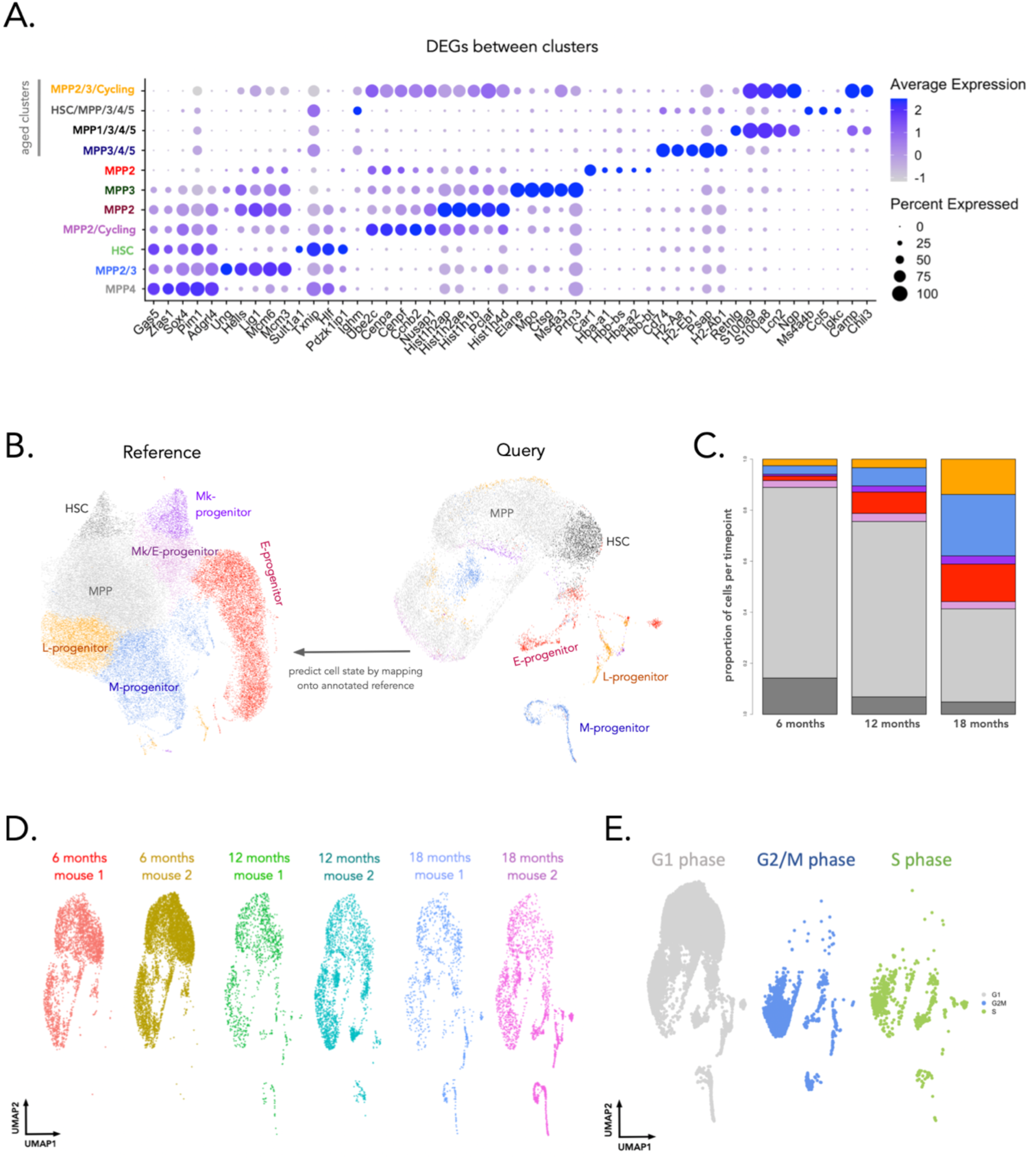
Supporting analyses for scRNAseq profiling of aged HSPCs. (A) Top 5 differentially expressed genes for each cluster. Differential gene expression analysis was performed using a logistic regression test as implemented in the Seurat R package and Bonferroni correction was applied to account for multiple testing. (B) Label transfer for supervised annotation of cell state. scRNAseq atlas of hematopoiesis (left – reference dataset) comprises 44,802 c-Kit+ and c-Kit+ Sca1+ hematopoietic progenitors^30^. Cell clustering and supervised assignment of cluster identity on this reference atlas were taken from ^31^. scRNAseq data from our study (query dataset) was then mapped onto this dataset using Seurat’s FindTransferAnchors and TransferData methods. (C) Barplot showing the relative proportion of cell-state definitions obtained by label transfer mapping. (D) Cells from each individual mouse overlaid onto the UMAP embedding of the integrated data. (E) Cells from different cell cycle stages overlaid onto the UMAP embedding of the integrated data. Cell cycle stage was annotated using the classifier-based approach from ^32^ implemented as the cyclone method in the scran R package.

**Figure S10:**
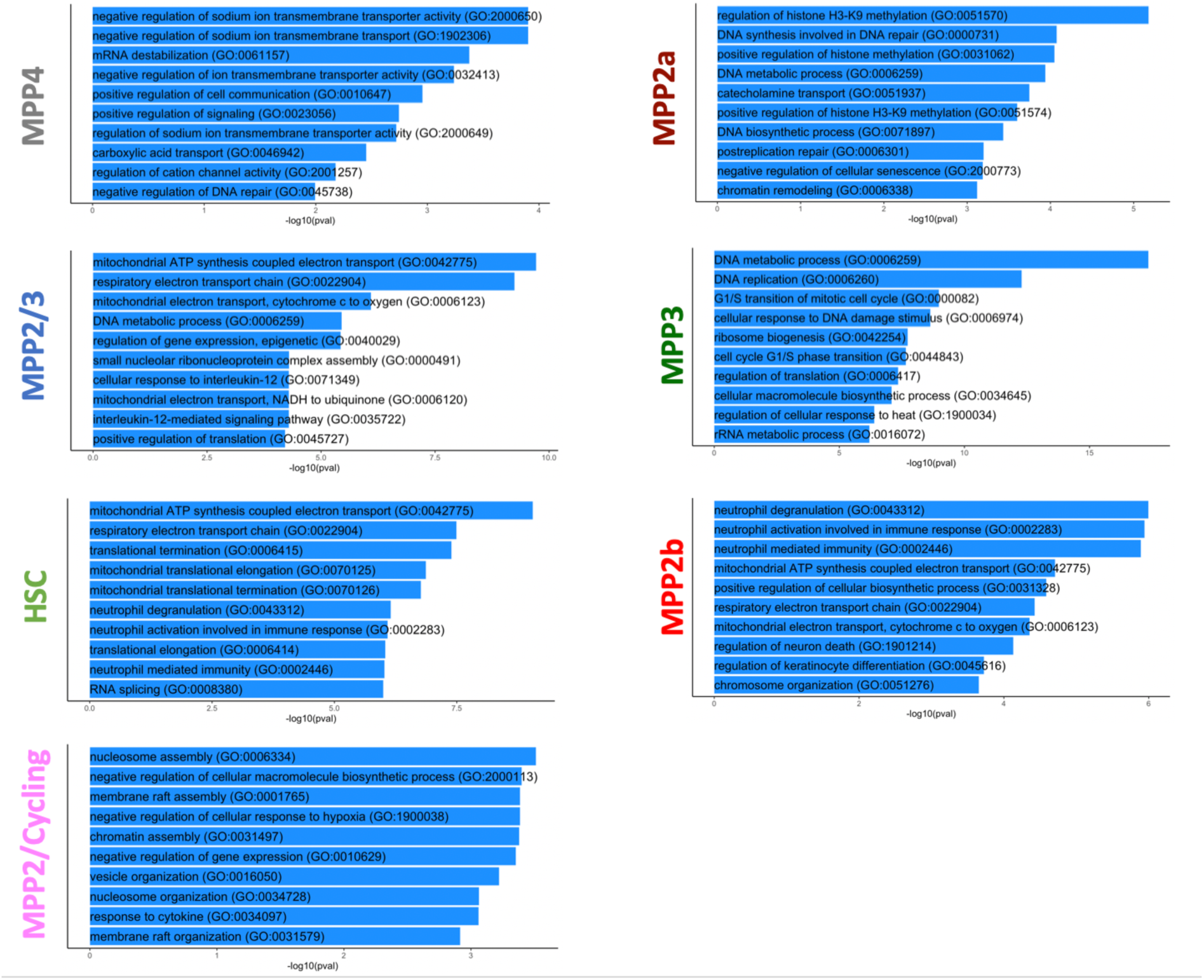
Pathways upregulated in HSPCs from aged mice across each cluster. Pathways enriched in HSPC clusters from mice aged 19 months compared to equivalent HSPC clusters from mice aged 6.5 months. Some clusters did not contain cells from 6.5 month old mice and were hence excluded from this analysis. Pathway analysis was performed using the enrichR R package using a variation of Fisher’s exact test, which also considers the size of each gene set when assessing its statistical significance.

**Figure S11:**
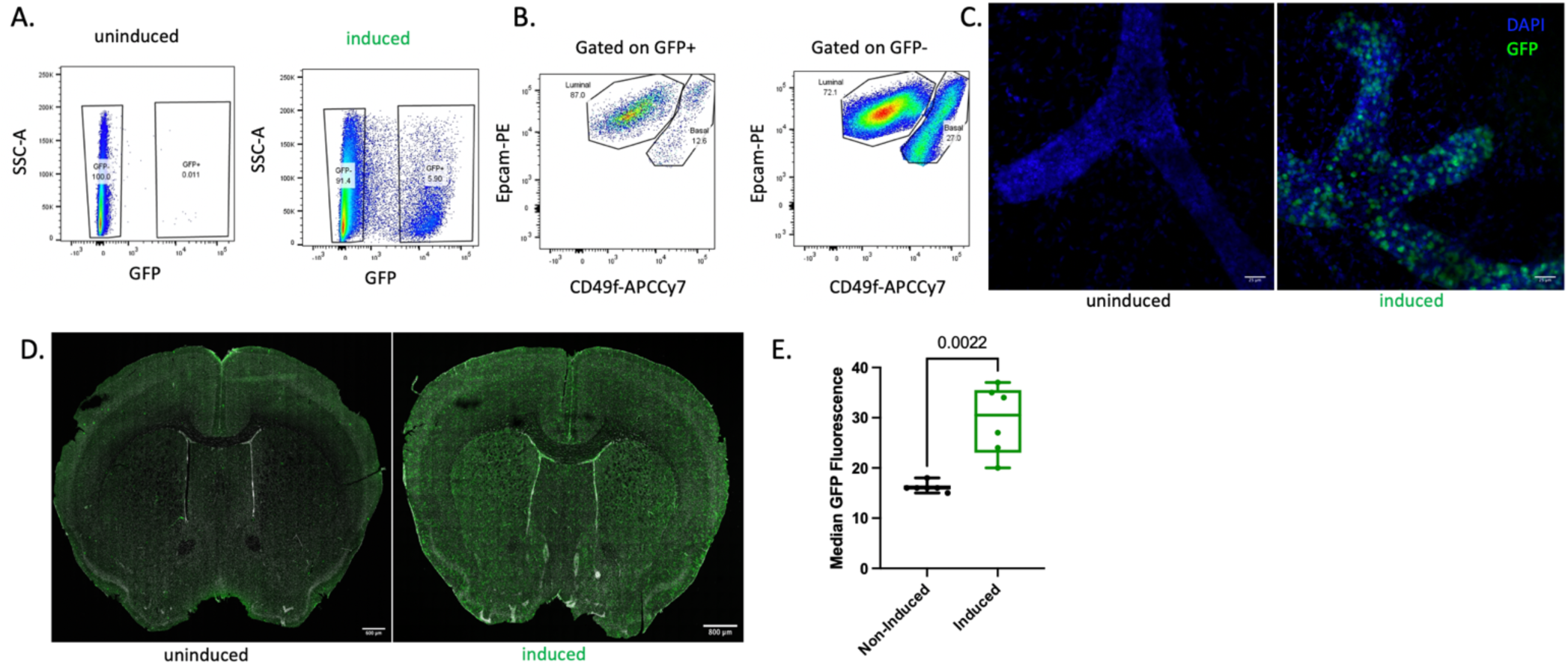
DRAG barcoding in the mammary gland and the brain. (A) Representative GFP+ cell population in mammary epithelial cells of an uninduced mouse (left) and induced mouse (right) 1 month post-induction. (B) Representative flow cytometry dot plots of luminal and basal cells gated within the mammary epithelial GFP positive (left) and GFP negative (right) populations. (C) Maximum projection of whole mount mammary gland from uninduced and induced DRAG mice 1 month post-induction, showing DAPI and GFP signal. (D) Exemplary images of GFP^+^ signal from barcoded cells in brain tissue sections from uninduced and induced DRAG mice and (E) Quantification of GFP fluorescence intensities in uninduced vs induced DRAG mice from different sections of the brain. Each point represents a different tissue section from 2 induced and 2 uninduced mice (3 tissue sections per mouse). Statistical comparisons were made using a Mann-Whitney test. Boxplots represent the median and IQR, whiskers extend to the min and max values.

**Figure S12.**
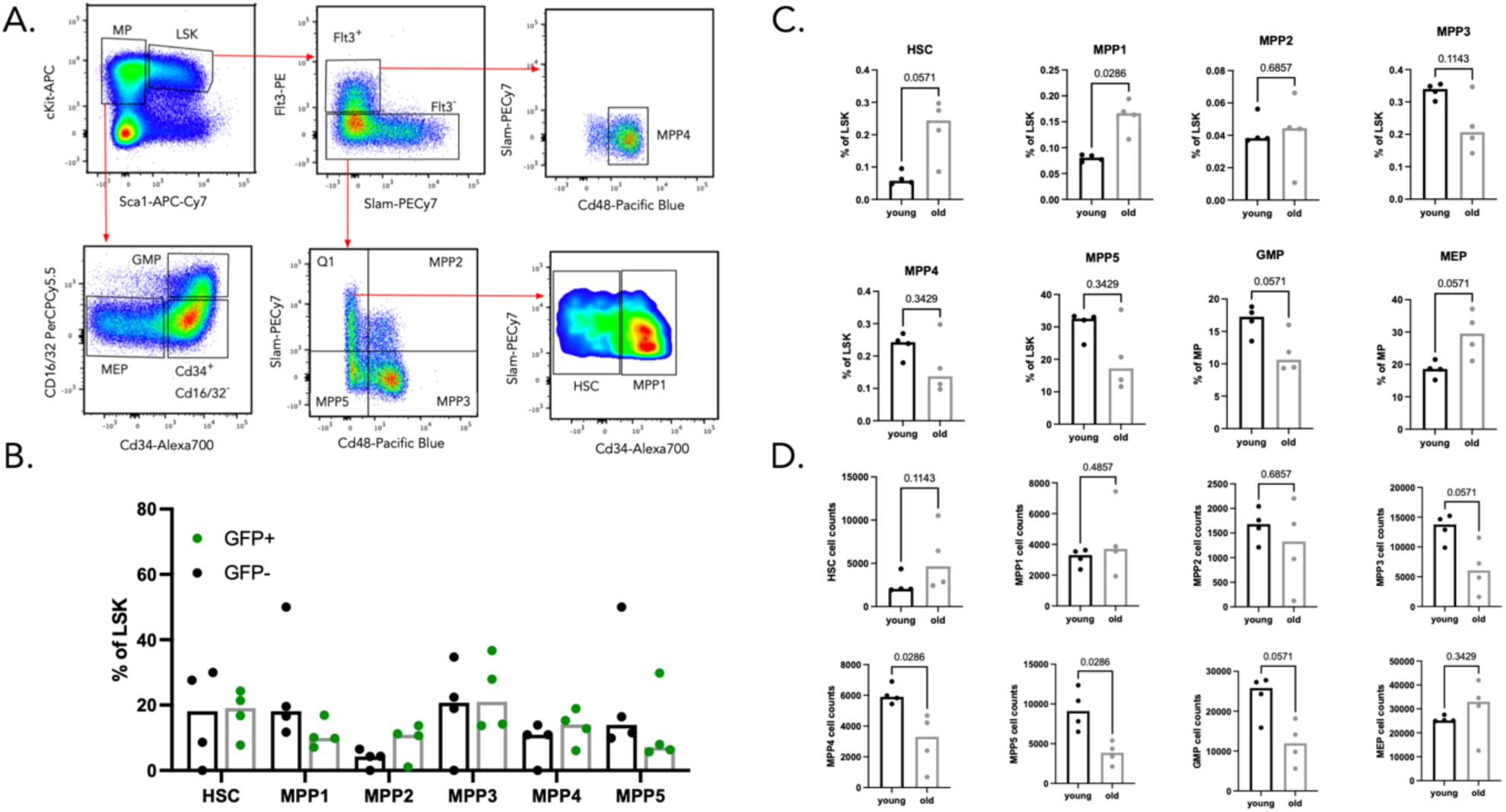
Flow cytometry analysis of HSPC and MP subsets in young and old mice. A. Gating strategy to identify the different HSPC and MP subsets. B. Flow cytometric quantification of the proportion of GFP+ and GFP− HSPC subsets in mice aged 19 months. Each point represents 1 mouse and n = 4. No statistically significant differences between GFP− and GFP+ representation amongst the HSPC subsets were observed. Statistical significance was tested using a paired Wilcoxon-Test. C. Quantification of HSPCs and MP subset frequencies between young (6.5 months) and old (19 months) mice. Each point represents 1 mouse and n = 4 mice. The Y-axis represents the percentage of each celltype amongst the entire cKit+ Sca1+ LSK compartment or the cKit+ Sca1-myeloid progenitor compartment. Statistical comparisons were made using a Mann-Whitney test. D. Quantification of HSPCs and MP subset cell counts between young (6.5 months) and old (19 months) mice. Each point represents 1 mouse and n = 4 mice. Statistical comparisons were made using a Mann-Whitney test.

